# Transcriptomic profiles of stress susceptibility and resilience in the amygdala and hippocampus

**DOI:** 10.1101/2023.02.08.527777

**Authors:** Kimberly L. P. Long, Sandra E. Muroy, Siamak K. Sorooshyari, Mee Jung Ko, Yanabah Jaques, Peter Sudmant, Daniela Kaufer

**Author notes:** These authors contributed equally to this work (Kimberly L. P. Long, Sandra E. Muroy). Department of Psychiatry and Behavioral Sciences, University of California, San Francisco, San Francisco, CA 94143, USA.

## Abstract

A single, severe episode of stress can bring about myriad responses amongst individuals, ranging from cognitive enhancement to debilitating and persistent anxiety; however, the biological mechanisms that contribute to resilience versus susceptibility to stress are poorly understood. The dentate gyrus (DG) of the hippocampus and the basolateral nucleus of the amygdala (BLA) are key limbic regions that are susceptible to the neural and hormonal effects of stress. Previous work has also shown that these regions contribute to individual variability in stress responses; however, the molecular mechanisms underlying the role of these regions in susceptibility and resilience are unknown. In this study, we profiled the transcriptomic signatures of the DG and BLA of rats with divergent behavioral outcomes after a single, severe stressor. We subjected rats to three hours of immobilization with exposure to fox urine and conducted a behavioral battery one week after stress to identify animals that showed persistent, high anxiety-like behavior. We then conducted bulk RNA sequencing of the DG and BLA from susceptible, resilient, and unexposed control rats. Differential gene expression analyses revealed that the molecular signatures separating each of the three groups were distinct and non-overlapping between the DG and BLA. In the amygdala, key genes associated with insulin and hormonal signaling corresponded with vulnerability. Specifically, *Inhbb, Rab31*, and *Ncoa3* were upregulated in the amygdala of stress-susceptible animals compared to resilient animals. In the hippocampus, increased expression of *Cartpt -* which encodes a key neuropeptide involved in reward, reinforcement, and stress responses - was strongly correlated with vulnerability to anxiety-like behavior. However, few other genes distinguished stress-susceptible animals from control animals, while a larger number of genes separated stress-resilient animals from control and stress-susceptible animals. Of these, *Rnf112, Tbx19*, and *UBALD1* distinguished resilient animals from both control and susceptible animals and were downregulated in resilience, suggesting that an active molecular response in the hippocampus facilitates protection from the long-term consequences of severe stress. These results provide novel insight into the mechanisms that bring about individual variability in the behavioral responses to stress and provide new targets for the advancement of therapies for stress-induced neuropsychiatric disorders.

## Introduction

Traumatic experiences bring about diverse cognitive responses in humans and animals, ranging along a continuous spectrum from enhanced performance to cognitive impairment^1^. In particular, a subset of individuals who experience trauma will develop long-lasting mood and anxiety disorders, such as posttraumatic stress disorder (PTSD)^2^, while others may exhibit beneficial outcomes, such as posttraumatic growth (PTG)^3^. Understanding the biological basis of individual variability may aid in developing new treatments to promote resilience; however, the neural and molecular mechanisms that bring about the various behavioral responses to trauma remain largely unknown.

The amygdala and hippocampus are key contributors in the brain to the physiological, affective, and behavioral responses to stress^4,5^. The amygdala regulates innate and learned emotional and behavioral responses to fear-inducing stimuli^6^, while the hippocampus contributes to context-dependent fear memory and generalized anxiety^7,8^. In addition, the hippocampus provides negative feedback to hypothalamic circuits governing the physiological stress response^9^. The amygdala and hippocampus are in turn regulated by stress^10^. Acute stress induces immediate changes to gene expression in each of these regions^11,12^, and such transcriptional changes can be long-lasting, particularly following chronic or traumatic stress^13–15^. The dentate gyrus (DG) of the hippocampus is particularly sensitive to acute and chronic stressors^16–18^, and we have previously found that this region displays cellular profiles that correlate with and contribute to behavioral outcomes after stress^19^. Similarly, acute and chronic stress induces structural and functional changes within the amygdala that are associated with increased anxiety-like behavior^20,21^. Structural and functional changes are often mediated by alterations in the transcriptional landscape. Ultimately, the transcriptional profiles of the hippocampus and amygdala may influence fear and anxiety behavior following stress and contribute to the susceptibility or resilience of an individual to a traumatic event^14,22^. We therefore hypothesized that transcriptional signatures in the amygdala and hippocampus would correlate with individual profiles of resilience or susceptibility to an acute stressor.

In this study, we investigated the transcriptional signatures of the amygdala and hippocampal dentate gyrus in rats with divergent behavioral outcomes after stress. We first subjected rats to a single, severe stress paradigm and conducted a behavioral battery to characterize long-term avoidance and startle behaviors. We selected stress-resilient and stress-vulnerable animals and collected tissue samples from the dentate gyrus and amygdala for transcriptomic analyses. Our results revealed that the broader transcriptome was not different between groups; however, we identified that select, region-specific sets of genes related to insulin signaling, neural plasticity, and stress corresponded with resilience and vulnerability after severe stress exposure.

## Methods

### Study design

The aim of this study was to determine amygdalar and hippocampal transcriptomic regulation of resilience and susceptibility to trauma. Rodents were exposed to an acute, severe stressor, and outcomes were assessed using physiological (serum corticosterone), behavioral (avoidance and startle assays), and transcriptional (RNA sequencing) measures. The experimental designs and methodology of outcome measures were chosen prior to initiating each experiment. For all experiments, animals were randomly assigned to experimental groups. Investigators were blind to stress conditions during behavioral assays and data quantification. The criteria for animal exclusion were experimental error and poor RNA quality (detailed below). Sample sizes and statistics for each experiment are provided in figure legends.

### Animals

All animal care and procedures were approved by the UC Berkeley Institutional Animal Care and Use Committee. A total of 40 adult (P65) male Sprague Dawley rats were purchased from Charles River and pair-housed on a 12:12 light-dark cycle (lights on at 0700 hours) in our facility at the University of California, Berkeley. Rats had *ad libitum* access to food and water and were given one week to acclimate to the facility before testing began. Rats underwent gentle handling (being picked up and held for about 2 minutes) for 5 days prior to stress.

### Stress

To study acute, severe trauma exposure, we coupled a single exposure to predator scent (fox urine) with 3 hours of immobilization in adult, male Sprague-Dawley rats. Rats were randomly assigned to undergo stress or remain in the home cage. Rats were run in pairs in cohorts of 10 animals each, with numbers of control and stress animals counterbalanced between cohorts. In the stress group (n = 20), rats underwent acute immobilization stress with exposure to predator scent. Cage mates were placed side-by-side in an empty cage inside a fume hood from 0900-1200 hours while being restrained in Decapicone bags (Braintree Scientific, Braintree, MA). A cotton ball infused with 1 mL of fox urine (Trap Shack Co. Red Fox Urine, amazon.com) was placed in the center of the cage. Blood sampling occurred throughout stress (detailed below). After cessation of stress, animals were released into a clean cage to allow for self-grooming. After 1 hour, rats were returned to a clean home cage. Rats in the control group (n = 20) received a cage change at the same time of day. To prevent transfer of stress and fox odors to control animals in the colony room, stress-exposed animals were housed in a separate room for 2 nights after stress before being returned to the colony.

### Weights

Animals were weighed on days -3 (handling), 0, +1, +2, +3, +7 (after behavior profiling day 1), +8 (after behavior profiling day 2), and +9 (day of sacrifice) relative to stress. One cohort of animals was mistakenly not weighed on the day of and day after stress; hence, these values are absent from plots of weight loss. Percent weight loss after stress was calculated as 100*(weight_Day +1_ - weight_Day 0_)/weight_Day 0_.

### Serum corticosterone sampling

To quantify hormonal stress responses, tail vein blood was collected from each rat for corticosterone sampling at 0 minutes, 30 minutes, and 3 hours into acute immobilization stress. Blood samples were centrifuged at 9,391 g for 20 minutes at 4°C, and serum was extracted and stored at -80°C. Samples from the 30-minute and 3-hour time points were assayed using a Corticosterone EIA kit (Arbor Assays, Ann Arbor, MI).

### Behavioral battery

Animals underwent a behavioral battery 7 days after stress exposure to characterize the extent of persistent behavioral changes. All behaviors were conducted between 0800-1400 hours. Prior to all tests, animals were brought to the testing space and allowed at least 30 min to acclimate. One day prior to stress, all animals went through a 5 min baseline open field test (OFT) under dim lighting (15 lux). Seven days after acute immobilization stress, all rats were individually profiled for anxiety-like behaviors using 6 different behavioral tests: OFT in a brightly lit environment (OFT Light), OFT in a dimly lit environment (OFT Dim), light/dark box (LD), elevated plus maze (EPM) under bright white light, EPM under dim red light, and acoustic startle response (ASR). Low light versions of the OFT and EPM were utilized as low anxiogenic versions of their full light counterparts, allowing for further characterization of exploratory and anxiety-like behavior. The battery spanned two days with the following sequence: Day 1 -- OFT Light, EPM Light, ASR; Day 2 – OFT Dim, LD, EPM Dim. After placing an animal into an arena, the experimenter exited the room. Animals were given 10 minutes of rest in the home cage in between each test. One control animal was excluded from subsequent analyses due to experimental error during behavior (animal was not given time to habituate to the room prior to behavior testing).

### Open field test (OFT)

Each rat was placed in an unenclosed plastic box (50 l x 50 w x 58 h in cm) and was given 10 minutes to freely explore the arena. All animals were placed along a wall at the beginning of the test. Behavior was recorded with cameras positioned above the arena and connected to GeoVision software (GeoVision Inc., Taiwan). Behavior was scored as latency to, frequency, and total amount of time spent in the center of the box (designated by a 25 × 25 cm square), as well as total distance traveled in the arena and in the center using EthoVision software (Noldus, Leesburg, VA). The OFT Light was conducted under full lighting (280 lux). The OFT Dim was conducted in a different but identically structured box in the same room under 15 lux. The arena was cleaned with 1% acetic acid followed by Formula 409 All Purpose Cleaner after each animal.

### Elevated plus maze (EPM)

Rats were allowed to explore an EPM for 10 min (arm dimensions: 10 w x 60 l in cm; closed arm enclosed by walls 51 cm in height; apparatus elevated 50 cm off the ground). The arms of the EPM were 10 cm wide. Behavior was recorded by a JVC Everio camera (JVCKENWOOD, Tokyo, Japan) mounted above the apparatus. The criteria for open arm exploration was considered as more than half of the body (and both forepaws) placed into the open arm. Latency to and total time spent in the exposed open arms, as well as time spent in the protected closed arms, were quantified by observers blind to condition. The EPM Light was conducted at 240 lux, while the EPM Dim was conducted under dim red light. The apparatus was cleaned with 1% Process NPD Disinfectant (STERIS Life Sciences) after each animal.

### Light-dark box (LD)

Each rat was placed in a structure consisting of an enclosed dark box separated by a divider with a small door leading to an unenclosed light box (each box 15 w x 15 l x 8 h in inches). All animals were placed into the dark half of the box and given 10 min to explore. Behavior was recorded by a JVC Everio camera (JVCKENWOOD, Tokyo, Japan) mounted above the apparatus. Measures for distance traveled as well as latency to, frequency, and total time spent in the exposed side were quantified by observers blind to condition. The arena was cleaned with 70% ethanol after each animal.

### Acoustic startle response (ASR)

Each rat was placed into an isolated Coulbourn sound-attenuating fear conditioning chamber (12 w x 10 l x 12 h in inches) and was exposed to 5 min of background noise (∼55 dB). This was followed by 70-110 dB white noise pulses lasting 10 ms, with an inter-stimulus interval of 15-30 s. All tones were calibrated with a handheld decibel meter each day prior to testing. Behavior was recorded over two different trials: habituation (110 dB tones presented 15 times to assess initial responses and subsequent habituation) and threshold determination (70-110 dB tones presented in pseudo-random order, with each tone played 5 times in total). Behavior was recorded using a Coulbourn Instruments camera connected to a computer with FreezeFrame software (Coulbourn Instruments, Whitehall, PA). The boxes were cleaned with 70% ethanol after each animal. Fear and startle behavior were assessed by Ethovision software analysis of activity change (measured as percent pixel change from frame to frame). The amplitude of startle was quantified as the maximum activity minus baseline activity in the 50 ms surrounding the startle pulse. Mean startle response was calculated as the mean of all startle amplitude scores across all 15 stimuli from the habituation phase. Sensitization was calculated as 100*[(mean startle amplitude to stimuli 13-15)-(mean startle amplitude to stimuli 1-3)]/(mean startle amplitude to stimuli 1-3).

### Composite scoring

To standardize and quantify behavior across multiple behavior tests, we adapted the method of Cutoff Behavioral Criteria developed by Cohen and Zohar^23^ and described in Long et al., 2021^19^. For each measure from the avoidance tests (OFTs, EPMs, LD) a behavioral cutoff criterion was defined by the 20^th^ percentile of the control distribution. For measures in which greater scores indicate greater anxiety-like behavior (latency to the anxiogenic zone, time spent in an anxiolytic zone), the 80^th^ percentile of the control group was used. Binary scoring was applied: Animals falling outside the criterion were marked as “affected” and received a score of 1 for that measure. Scores were then summed across all tests (minimum 0, maximum 20). High scores represent consistent anxiety-like behavior across all tests.

### Fresh tissue collection and RNA Sequencing

On day 9 after stress, animals were anesthetized with isoflurane and rapidly decapitated. Brains were extracted, flash frozen by floating on liquid nitrogen, and stored at -80°C. A subset of 12 animals were selected for RNA sequencing based on composite anxiety-like behavior scores: 3 stress-exposed animals with low scores (0-2), 3 stress-exposed animals with high scores (13-15), and 6 control animals from across the spectrum (0-6, average = 3.67). Sections of 200 μm thickness were collected under sterile conditions on a Leica cryostat, and punches of 0.75 mm diameter were taken from the dorsal dentate gyrus and basolateral amygdala (rat AP coordinates -2.9 to -5.28 relative to Bregma) using a Rapid-Core Sampling Tool (Electron Microscopy Sciences, Hatfield, PA). Punches were separated by hemisphere, and 7-9 punches were collected per hemisphere. Samples were stored at -80°C until processing. Tissue punches from the left hemisphere were homogenized in TRIzol reagent (Invitrogen, Waltham, MA), and RNA was extracted and treated with DNase (DNase I, RNase-free, New England Biolabs, Ipswich, MA) according to the TRIzol reagent user guide (Pub. No. MAN0001271, Rev. A.0). For RNA sequencing, all postprocessing (including cDNA library preparation) and sequencing was performed by the Vincent J. Coates Genomics Sequencing Laboratory at UC Berkeley. RNA quality scores were determined for each sample on an Agilent 2100 Bioanalyzer (Agilent Technologies, Santa Clara, CA). One sample (from a control animal with composite behavior score of 0) had low RNA quality, and RNA was re-extracted but later excluded from analysis (see Statistical Analysis section). RNA was poly-A selected, and library preparation was conducted on a Biomek FXp with Kapa Biosystems reagents. Sequencing was performed on the Illumina HiSeq4000 (Illumina, San Diego, CA).

Reads were aligned to the *Rattus norvegicus* Rn5 genome assembly using Spliced Transcripts Alignment to a Reference aligner (STAR/2.6.0a)^24^ with gene annotations provided from the ensemble build 75. Transcript/gene level quantification and differential gene expression were performed using rsem/1.3.1 + ebseq^25,26^. Gene Set Enrichment Analysis (GSEA)^27,28^ was performed on sets of differentially expressed genes (FDR < 0.05) to determine enriched biological themes.

### Statistical Analysis

All data are presented as mean ± standard error of the mean (SEM). Two sample comparisons were performed by two-tailed student’s t-test. To compare the relationships between behavioral and transcriptional measures, we computed Pearson correlations. In all tests, the alpha value was set at 0.05. Analyses were performed using IBM SPSS 19 (SPSS, Inc., Chicago, IL) and GraphPad Prism version 9.1.1 for Mac OS X (GraphPad Software, San Diego, California USA, www.graphpad.com). Hierarchical clustering of behavior was performed with the Seaborn package in Python (Version 3.7.1, Python Software Foundation, Wilmington, DE). For RNA-seq data, the R EBSeq^26^ package’s posterior probability of being differentially expressed (PPDE > 0.95, FDR <0.05) was used to determine statistical significance. One animal was excluded from all analyses due to experimental error during behavior assays (see Behavioral battery section above), and one hippocampal RNA sample was excluded from analyses due to contamination. Specifically, the sample was flagged for low RNA quality and was subsequently re-extracted from the remaining homogenized tissue. Although the re-extracted sample had a passing RNA integrity number, principal component analysis revealed that the sample did not cluster with any other samples (Fig. S1), suggesting that re-extraction compromised RNA integrity or introduced contamination. As a result, this sample was excluded from further analyses. Similarly, one amygdala sample was excluded from analysis as it did not cluster in PC space with any other samples.

## Results

### Acute, severe stress induces a potent stress response and a range of behavioral outcomes

To model acute trauma exposure, we subjected 20 adult, male Sprague-Dawley rats to immobilization with predator scent stress (fox urine). Another 19 animals served as controls and remained minimally disturbed in the home cage (Fig. 1A). We have previously demonstrated that this acute, combinatorial design is a severe stressor that elicits a potent physiological stress response in rats^19,29,30^. To identify subsets of animals displaying persistent high or low anxiety-like behavior after stress, all animals were allowed 7 days of recovery with minimal disturbance before undergoing a 2-day behavioral profiling battery. This battery consisted of multiple tests of approach-avoidance conflict and acoustic startle (Fig. 1A). Consistent with our previous work^19^, we found that stress-exposed rats displayed considerable inter-individual variability in each behavioral metric (Fig. S2) and that individuals displayed consistent behavioral trends across avoidance assays (Fig. S3). Composite behavioral scoring revealed that a subset of stress-exposed rats displayed considerable, persistent anxiety-like behavior (Fig. 1B). We selected 3 of these stress-susceptible animals and 3 stress-resilient animals from the opposite end of the spectrum for subsequent transcriptomic analyses (Fig. 1C-D). Physiologically, stress-exposed rats lost significantly more weight by the day after stress than control animals (Fig. 1E), and weight changes trended towards a negative correlation with composite anxiety-like behavior scores (Fig. 1F). Stress-exposed rats also displayed high circulating corticosterone at 30 minutes and 3 hours into stress (Fig. 1G). Corticosterone at 30 minutes was not correlated with behavioral scores (Fig. 1H). However, serum corticosterone at 3 hours (the end of stress) was significantly, positively correlated with overall composite scores and avoidance sub-scores (Fig. S4B), suggesting that sustained high release of corticosterone may contribute to subsequent anxiety-like behavior. Together, these data suggest that a single, severe stressor induces persistent avoidance behavior in a subset of rats that is correlated with sustained high corticosterone release during the stressor.

**Figure 1:**
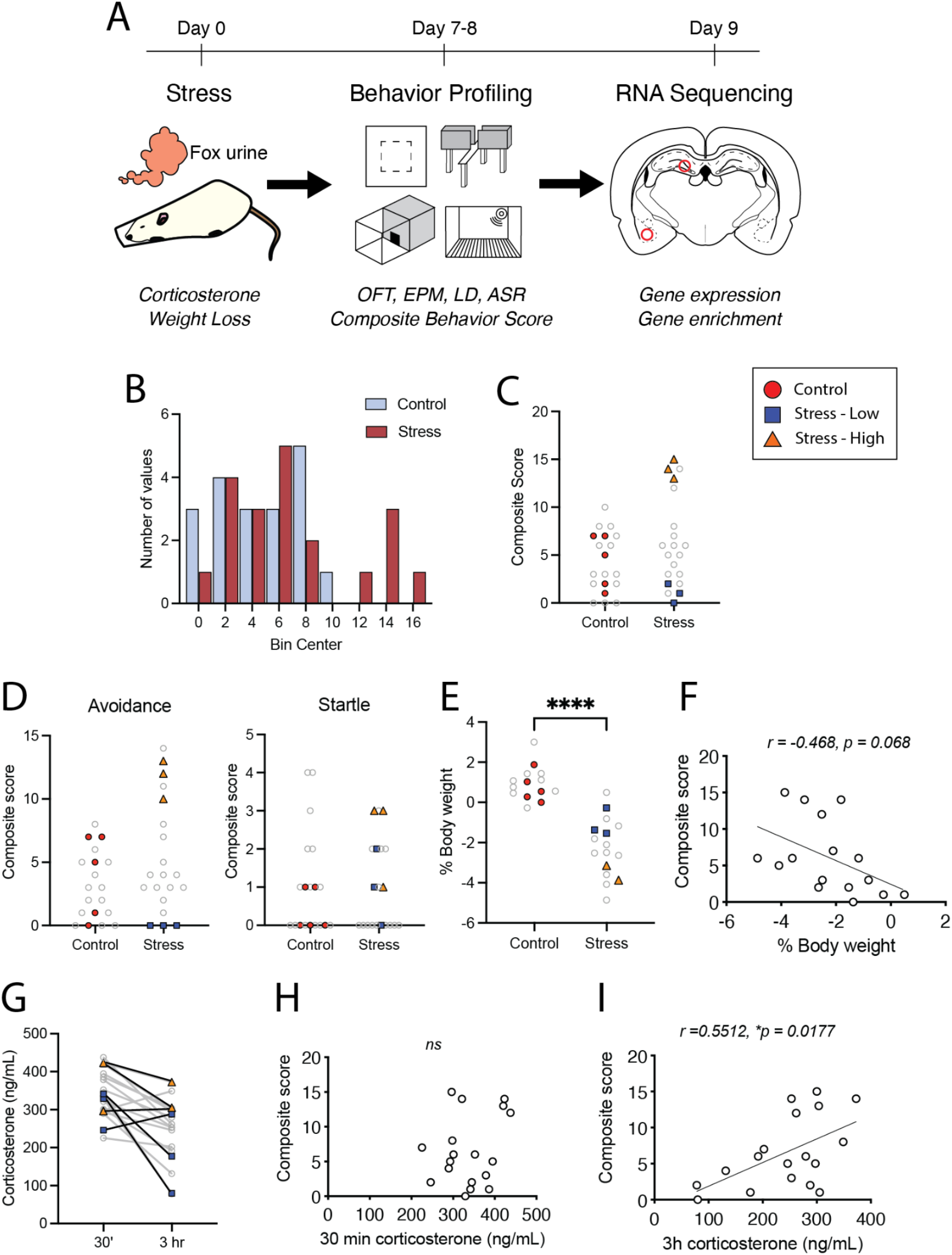
Experimental design and selection of control, stress-susceptible, and stress-resilient rats. A. Experimental design. Male Sprague-Dawley rats were subjected to a single, severe stressor (3 hours of immobilization with exposure to fox urine). Body weight was measured on the day of and day after stress, and serum samples were collected for corticosterone measurements at 30 minutes and 3 hours into stress. Behavior profiling occurred on days 7 and 8 after stress, and whole brains were flash frozen on liquid nitrogen on day 9 for tissue punch collections and bulk RNA sequencing from the dorsal dentate gyrus and basolateral amygdala. B. Histogram of composite anxiety-like behavior scores (bin size = 2). C. Distribution of individual composite behavior scores from control and stress-exposed rats. Highlighted points indicate the individual animals selected for RNA sequencing. D. Sub-scores of composite behavior derived from avoidance (left) and startle (right) assays. E. Percent changes in body weight from day 0 (day of stress) to day 1. Negative values indicate weight loss (2-tailed unpaired t test with Welch’s correction for unequal variance, t(24.38) = 7.466, p < 0.0001). F. Pearson correlation of percent body weight change to composite behavior scores (r = -0.468, p = 0.068). G. Serum corticosterone measurements at 30 minutes and 3 hours into stress exposure. H. Pearson correlation of corticosterone at 30 minutes to composite behavior scores (r = 0.2443, p = 0.3287). I. Pearson correlation of corticosterone at 3 hours to composite behavior scores (r = 0.5512, p = 0.0177). OFT, open field test. EPM, elevated plus maze. LD, light-dark box. ASR, acoustic startle response.

### The amygdala and dentate gyrus display distinct transcriptomic changes following acute, severe stress

Following behavior profiling, we conducted RNA-sequencing (RNA-seq) on bulk tissue isolated from the left hippocampal dentate gyrus (DG) and amygdalae (Fig. 1A). We selected stress-exposed rats with low (“stress-low”, n = 3) and high (“stress-high”, n = 3) composite anxiety-like scores as well as control animals (n = 5 DG, 6 amygdala, Fig. 1C, Fig. 2A). Of the selected animals, mean composite anxiety-like behavior scores were 4.4 ± 1.2 for control animals, 1.0 ± 0.58 for stress-low animals, and 14.0 ± 0.58 for stress-high animals. Principal component analysis (PCA) of normalized gene expression values for both the amygdala and hippocampus DG revealed that rats did not clearly cluster by condition in PC space (Fig. S1A-D). However, pairwise comparisons between each of the 3 conditions yielded select sets of differentially expressed genes (DEGs; FDR <0.05) between our 3 conditions in both regions (Fig. 2B-C). There was no overlap between the sets of DEGs in the amygdala versus the DG (Fig. 2D), supporting the idea that distinct mechanisms underlie stress-induced structural and functional changes in these 2 regions.

**Figure 2:**
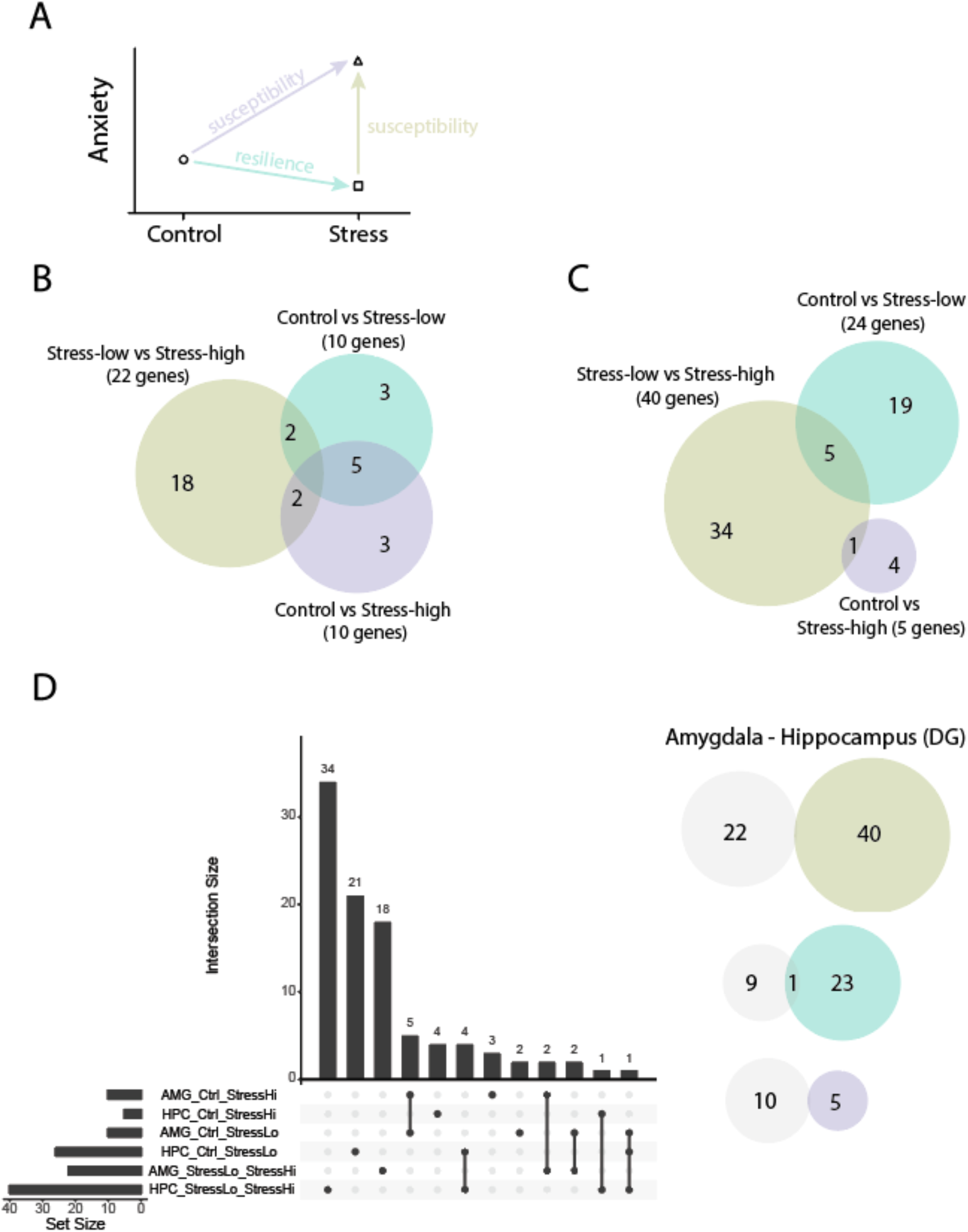
Regionally distinct transcriptomic changes in the amygdala versus dentate gyrus following acute, severe stress. A. Schematic for differential gene expression (DGE) analysis. Arrows denote pairwise comparisons yielding susceptibility or resilience-like genes. B. Venn diagram showing DGE results for the amygdala for our 3 pairwise comparisons. C. Venn diagram showing DGE analysis results for the hippocampus dentate gyrus (DG). D. Upset plot displaying overlaps between the 3 pairwise comparisons in the amygdala versus DG. Note that gene sets are largely independent with few to no overlaps between the 2 regions.

### Acute, severe stress affects expression of genes related to insulin and hormone signaling the in amygdala of susceptible and resilient animals

In the amygdala, the comparison between stress-low and stress-high animals yielded 22 DEGs (Fig. 3A). Of these, 8 genes were downregulated, while 14 genes were upregulated in stress-high animals. Between control and stress-low animals, there were 10 DEGs, 2 of which were downregulated in stress-low animals (Fig. 3B). The comparison between control and stress-high animals also yielded 10 DEGs, 3 of which were upregulated in stress-high animals (Fig. 3C). Two genes overlapped between the stress-low/stress-high and control/stress-low comparisons (*Gns* and *Rasl10b*), both of which were downregulated in stress-low animals in each comparison. Two genes overlapped between stress-low/stress-high and control/stress-high (*Rnf11l* and *CALCOCO1*), with *Rnf11l* downregulated in stress-high animals. Lastly, 5 genes overlapped between control/stress-low and control/stress-high comparisons (*Smg6, Cyp11b3, Mrgbp, MYH9*, and *TPI1*). *Smg6* was upregulated in stress-exposed animals, while DEG status of the remaining genes may have been driven by outlier values.

**Figure 3:**
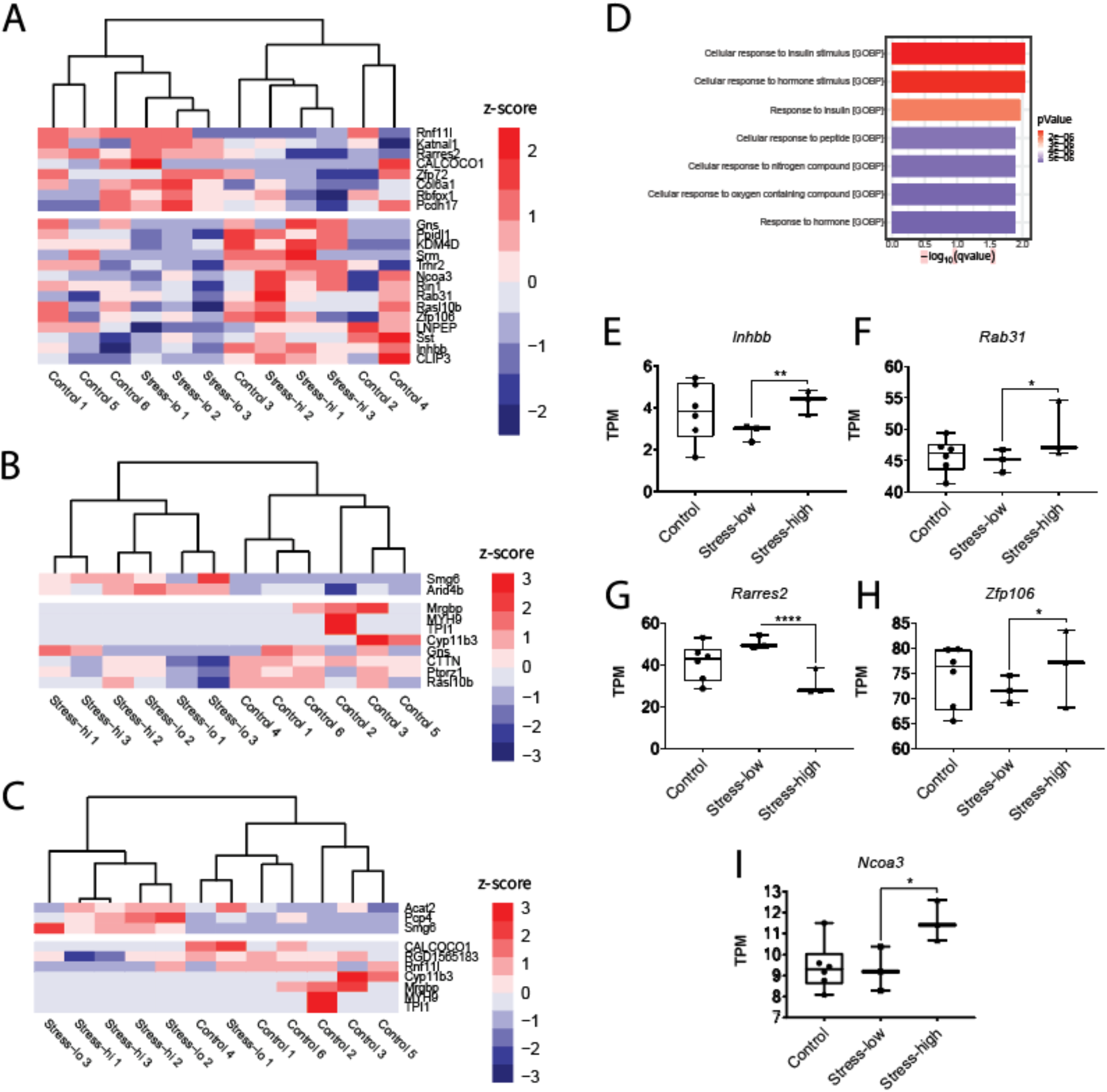
Gene expression changes in the amygdala of susceptible and resilient animals following acute, severe stress. A. Heatmap showing differentially expressed genes (DEGs) for the stress-low versus stress-high comparison. B. Heatmap of DEGs for the control versus stress-low comparison. C. Heatmap of DEGs for the control versus stress-high comparison. For all heatmaps: dendrograms show the clustering of samples (columns). Rows/genes were clustered based on Pearson correlation but explicit dendrograms were omitted for clarity of presentation. Breaks denote the separation between up- and downregulated genes. D. Results of Gene Set Enrichment Analysis (GSEA) showing the top significantly enriched Gene Ontology (GO) categories. E-I. Gene expression (in TPM) for genes comprising the top 2 GO Biological Process (GOBP) categories. Asterisks denote the FDR (i.e., 1-Posterior probability of being differentially expressed (PPDE)); * ≤ 0.05, ** ≤ 0.01, * ** ≤ 0.0001.

To understand the biological processes in the amygdala that might contribute to our observed anxiety-like phenotypes, we conducted Gene Set Enrichment Analysis (GSEA)^27,28^. We focused on the largest set of DEGs (between stress-low and stress-high animals: 22 genes) and found significant enrichment for gene ontology biological process (GOBP) categories including ‘cellular response to insulin stimulus’ and ‘cellular response to hormone stimulus’ (Fig. 3D). Individual transcripts per million (TPM) values for the top 2 GOBP category genes (*Inhibin beta B chain*/*Activin beta B chain [Inhbb], Rab31, member RAS oncogene family [Rab31*], *Retinoic acid receptor responder protein 2* [*Rarres2*], *Zinc finger protein 106* [*Zfp106*], and *Nuclear receptor coactivator 3* [*Ncoa3*]) are shown in Fig. 3 panels E-I. In addition to their role in response to insulin or hormone stimulus, 3 of the genes are also specifically associated with modulation of anxiety-like behaviors in mice: *Inhbb*^31–33^, *Rab31*^34^, and *Ncoa3*^35–37^. All 3 were significantly upregulated when comparing stress-high to the stress-low condition (Fig. 3E, F, I). Among the top 2 GOBP category genes, *Rarres2* (which plays a role in insulin sensitivity) showed significant correlations to composite anxiety-like behavior scores and avoidance sub-scores (Fig. S5).

### Increased Cartpt expression in the dentate gyrus correlates with susceptibility to anxiety-like behavior after acute, severe stress exposure

In the DG of the hippocampus, the comparison between stress-low and stress-high animals yielded 40 DEGs (Fig. 4A), 17 of which were downregulated and 23 upregulated in stress-low animals. Among control and stress-low animals, there were 24 DEGs (Fig. 4B). Of these, 9 were upregulated and 15 downregulated in stress-low animals. Between control and stress-high animals, there were only 5 DEGs, with 3 genes downregulated and 2 upregulated in stress-high animals (Fig. 4C). We examined the genetic overlap between each of the 3 pairwise comparisons (Fig. 3D). In comparing control/stress-low and stress-low/stress-high (‘resilience’ signature), we found 5 overlapping genes: *Rnf112* (*Ring finger protein 112; Zinc finger protein 179* [*ZNF179*]; *Neurolastin*), *Tbx19* (*T-box transcription factor 19*), *UBALD1* (*UBA Like Domain Containing 1*), *ATN1* (*Atrophin 1*), and *CTTN* (*Cortactin*). Individual TPM values for *Rnf112* (Fig. 4E), *Tbx19* (Fig. 4F), and *UBALD1* (Fig. 4G) indicate that expression of each of these genes is significantly downregulated in stress-low animals when compared to controls and stress-high animals, suggesting active gene expression changes are occurring in these ‘stress resilient’ animals. Differential expression of *ATN1* and *CTTN* was driven by outlier values (Fig. S6). The overlap between control/stress-high and stress-low/stress-high DEGs (‘susceptibility’ signature) revealed a single shared gene - *Cartpt* (cocaine and amphetamine regulated transcript [CART] prepropeptide). Transcription of *Cartpt* was significantly upregulated in the stress-high group (Fig. 4H), while stress-low values did not differ from controls. *Cartpt* expression was positively correlated to the composite anxiety-like behavior scores and avoidance sub-scores, but not startle sub-scores (Fig. S7).

**Figure 4:**
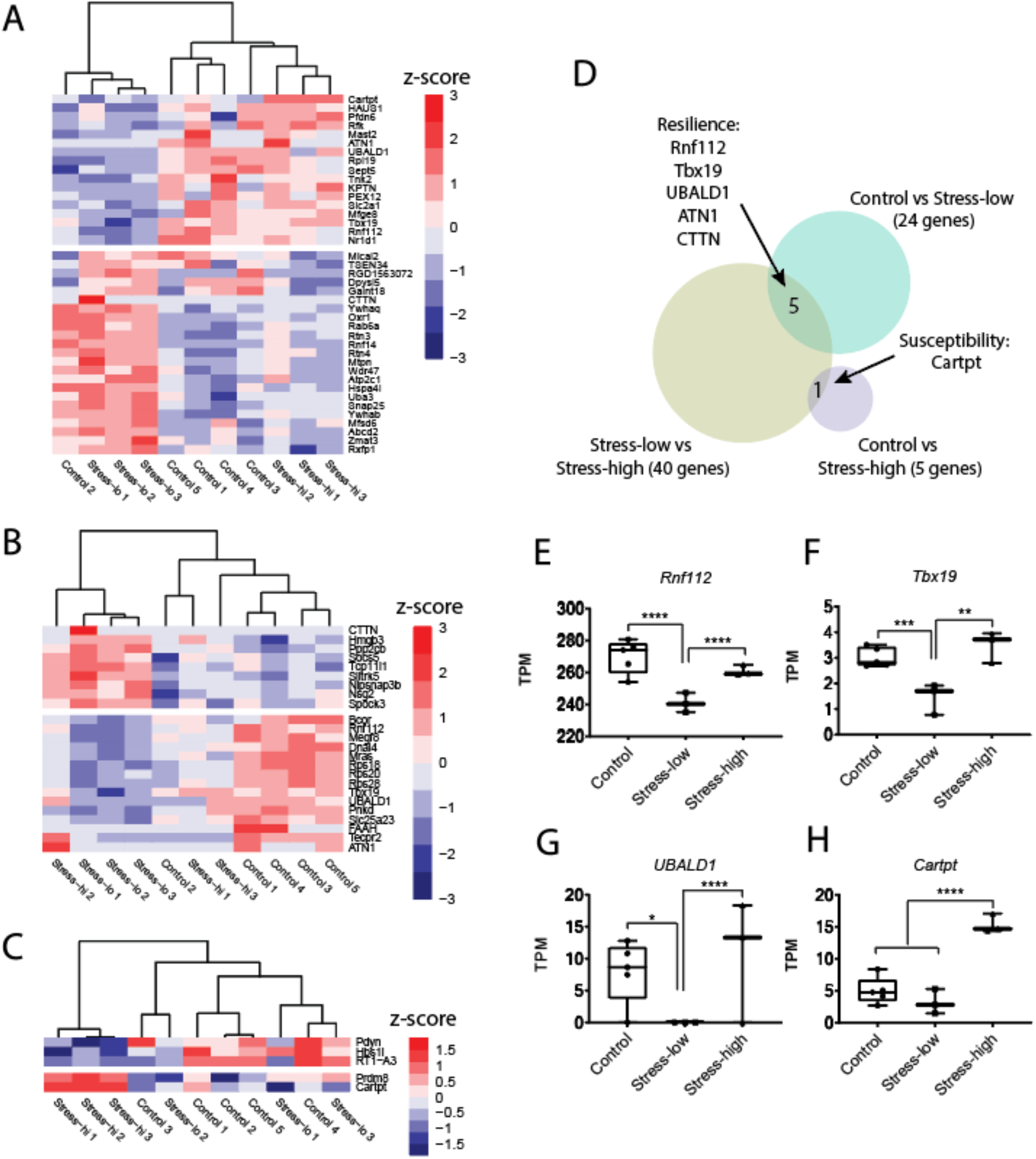
Gene expression changes in the hippocampal dentate gyrus of susceptible and resilient animals following acute, severe stress. A. Heatmap showing differentially expressed genes (DEGs) for the stress-low versus stress-high comparison. B. Heatmap of DEGs for the control versus stress-low comparison. C. Heatmap of DEGs for the control versus stress-high comparison. For all heatmaps: dendrograms show the clustering of samples (columns). Rows/genes were clustered based on Pearson correlation but explicit dendrograms were omitted. Breaks denote the separation between up- and downregulated genes. D. Venn diagram displaying genes in overlaps between: 1) the stress-low versus stress-high and control versus stress-low comparisons (‘Resilience’ signature genes) and 2) stress-low versus stress-high and control versus stress-high comparisons (‘Susceptibility’ signature gene). E-G. Gene expression (in TPM) for *Rnf112, Tbx19* and *UBALD1* is significantly downregulated in the stress-low group compared to stress-high and control animals. H. *Cartpt* gene expression (in TPM) is significantly upregulated in the stress-high group compared to both stress-low and control animals. Asterisks denote the FDR (i.e., 1-Posterior probability of being differentially expressed (PPDE)); * ≤ 0.05, ** ≤ 0.01, \*\*\**p* ≤ 0.001, **** ≤ 0.0001.

GSEA indicated that sets of DEGs in the DG for our 3 comparisons did not yield significant biological enrichment. Hence, we describe all individual DG DEGs and their currently known functions in Tables 1-3. Many of the genes identified in the stress-low vs stress-high comparison play roles in the regulation of dendritic spine structure and function, neurite outgrowth, anxiety, mood, and psychiatric disorders.

**Table 1.**
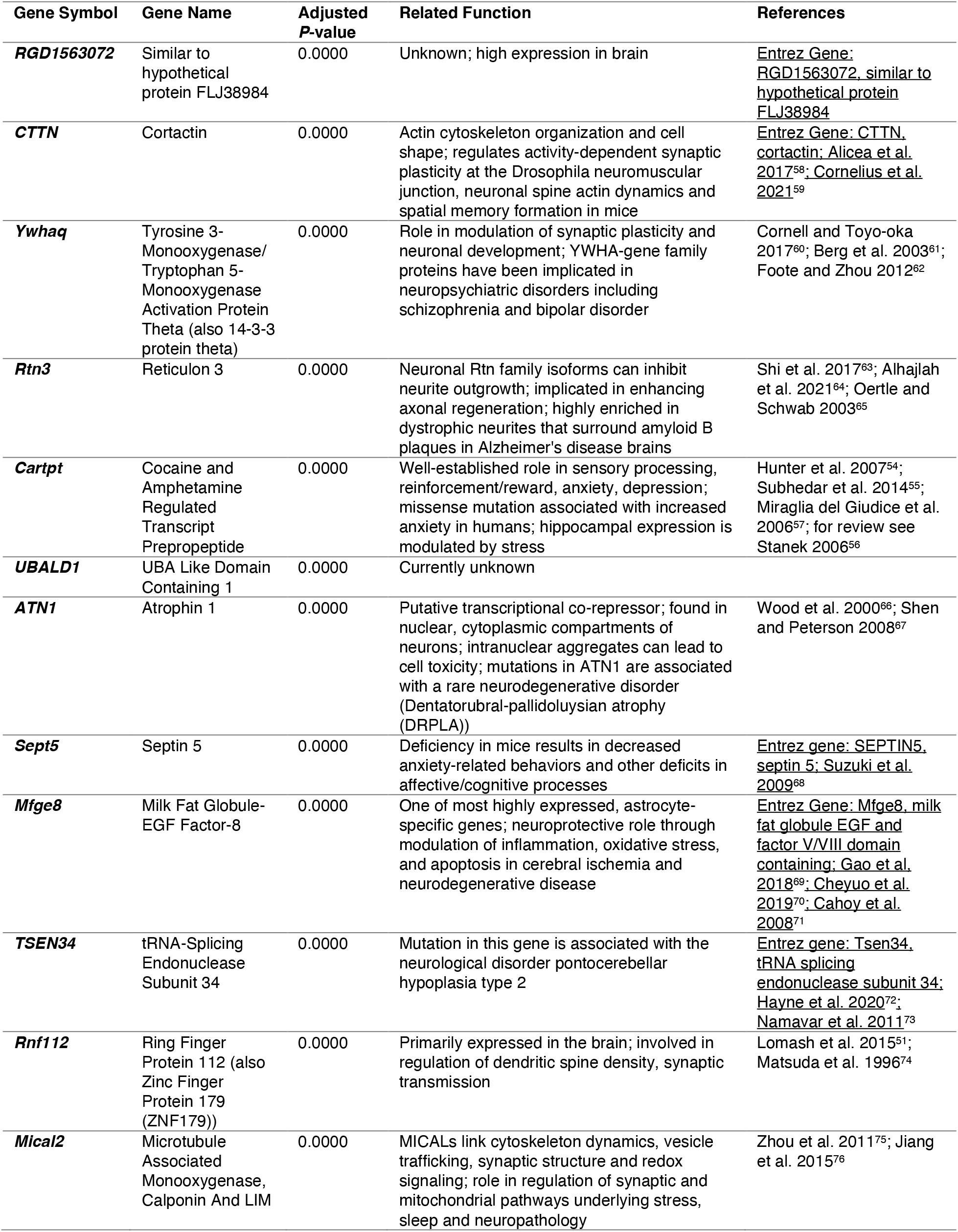
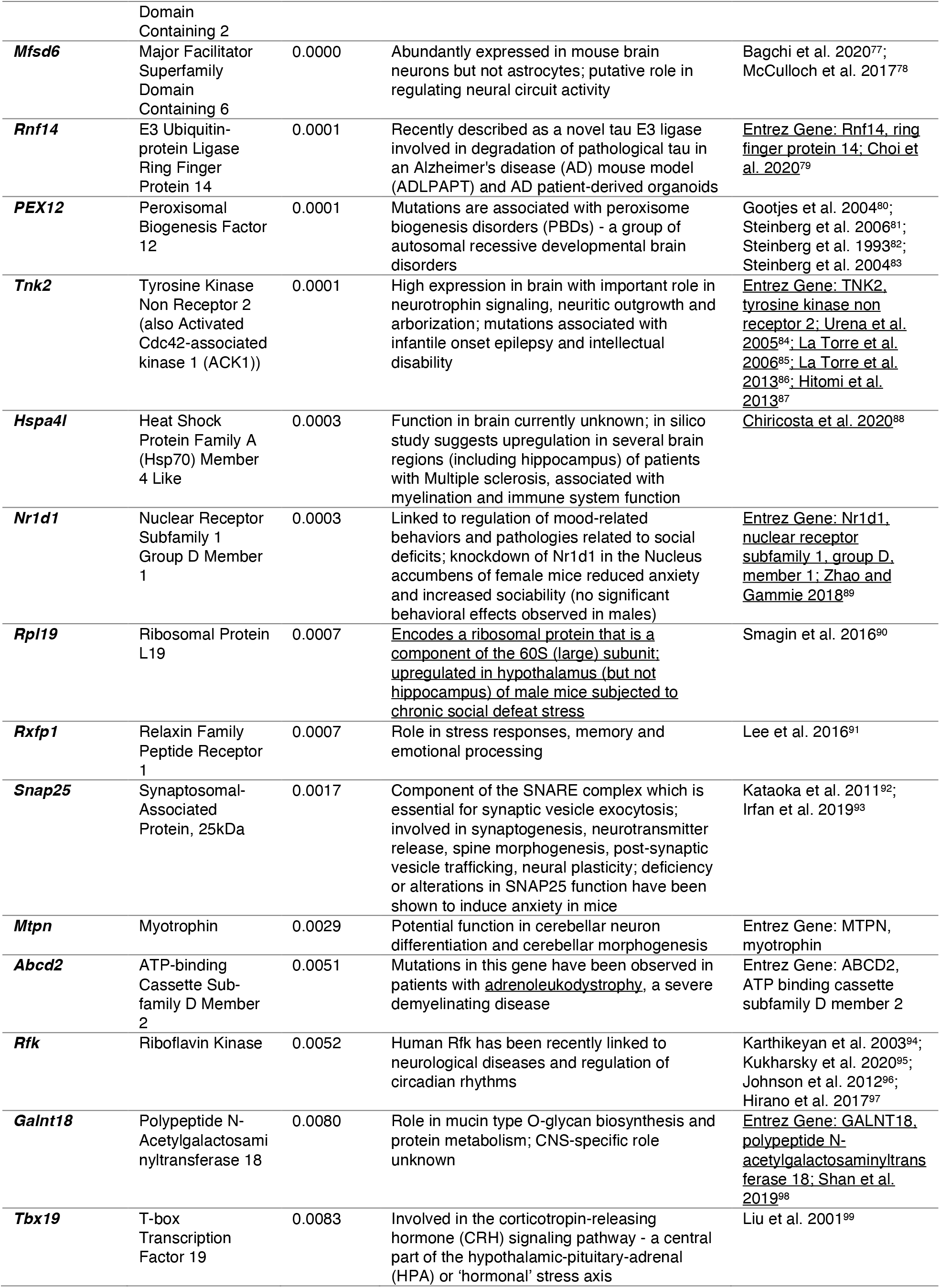
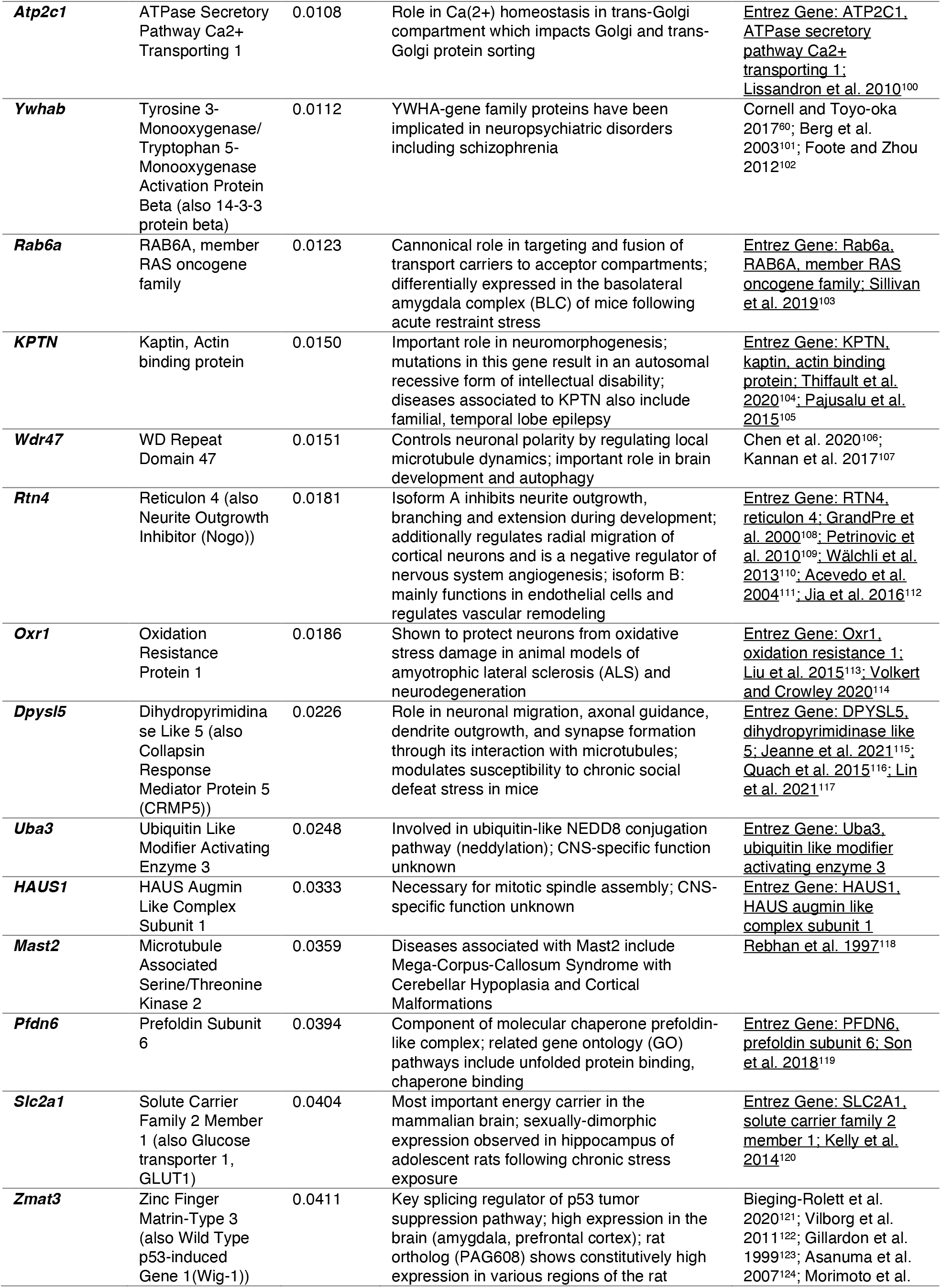
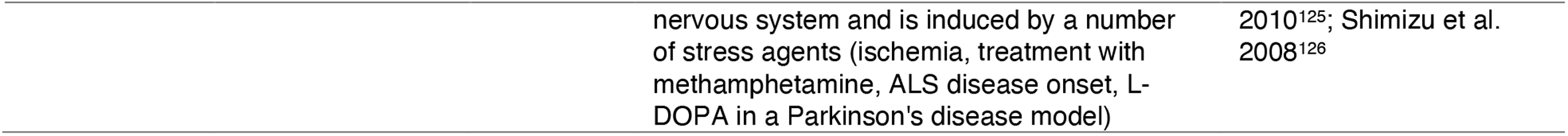
Differentially Expressed Genes Between Stress Low and Stress High Animals.

**Table 2.** Differentially Expressed Genes Between Stress Low and Control Animals.

| Gene Symbol | Gene Name | Adjusted P-value | Related Function | References |
| --- | --- | --- | --- | --- |
| ***CTTN*** | Cortactin | 0.0000 | Role in actin cytoskeleton organization, cell shape; regulates activity-dependent synaptic plasticity (*Drosophila* NMJ) and neuronal spine actin dynamics, spatial memory formation in mice | Entrez Gene: CTTN, cortactin; Alicea et al. 2017<sup>58</sup>; Cornelius et al. 2021<sup>59</sup> |
| ***Mras*** | Muscle RAS Oncogene Homolog | 0.0000 | Member of RAS family of small GTPases, role in signal transduction mediating cell growth and differentiation; knockout of M-Ras in mice resulted in deficits in social interaction, increased fear-related behavior | Entrez Gene: MRAS, muscle RAS oncogene; Ehrhardt et al. 2015<sup>127</sup> |
| ***Megf8*** | Multiple EGF Like Domains 8 | 0.0000 | Role in Bone Morphogenetic Protein 4 (BMP4) signaling in trigeminal ganglion neurons, important for axon guidance in the peripheral nervous system | Engelhard et al. 2013<sup>128</sup> |
| ***ATN1*** | Atrophin 1 | 0.0000 | Putative role as transcriptional co-repressor; expressed in neurons; intranuclear aggregates can lead to cell toxicity; ATN1 mutations are associated with a rare neurodegenerative disorder (Dentatorubral-pallidoluysian atrophy (DRPLA)) | Wood et al. 2000<sup>66</sup>; Shen and Peterson 2008<sup>67</sup> |
| ***Rnf112*** | Ring Finger Protein 112 (also Zinc Finger Protein 179 (ZNF179)) | 0.0000 | Member of the RING finger protein family of transcription factors; primary expression in brain; involved in regulation of dendritic spine density, synaptic transmission | Lomash et al. 2015<sup>51</sup>; Matsuda et al. 1996<sup>74</sup> |
| ***FAAH*** | Fatty Acid Amide Hydrolase | 0.0000 | Encodes protein involved in primary and secondary fatty acid amide hydrolysis (including neuromodulatory anandamide, oleamide); modulation of anandamide signaling (via FAAH) is a therapeutic target for stress-related neuropsychiatric diseases | Entrez Gene: FAAH, fatty acid amide hydrolase; Gunduz-Cinar et al. 2014<sup>129</sup>; Morena et al. 2019<sup>130</sup> |
| ***Spock3*** | SPARC/Osteonectin, Cwcv and Kazal-Like Domains Proteoglycan 3 | 0.0000 | Related pathways include extracellular matrix degradation and ERK signaling; putative risk gene for adult ADHD and personality disorders | Entrez Gene: SPOCK3, SPARC (osteonectin), cwcv and kazal like domains proteoglycan 3; Weber et al. 2014<sup>131</sup> |
| ***Rpl19*** | Ribosomal protein L9-like? | 0.0001 | Part of 60S (large) ribosomal subunit; one study found modulation of Rpl19 expression in hypothalamus following chronic social defeat in mice | Entrez Gene: RPL19, ribosomal protein L19; Smagin et al. 2015<sup>132</sup> |
| ***Pnkd*** | Paroxysmal Nonkinesigenic Dyskinesia (PNKD) Metallo-Beta-Lactamase Domain Containing | 0.0002 | Mutations are associated to the movement disorder paroxysmal non-kinesigenic dyskinesia (characterized by involuntary unilateral or bilateral movements); association with Tourette Disorder or Tic disorder | Erro 1993<sup>133</sup>; Sun et al. 2018<sup>134</sup> |
| ***Tcp11l1*** | T-Complex 11 Like 1 | 0.0002 | Currently unknown | Entrez Gene: TCP11L1, t-complex 11 like 1 |
| ***Tbx19*** | T-box Transcription Factor 19 | 0.0007 | Involved in corticotropin-releasing hormone (CRH) signaling pathway - a central part of the hypothalamic-pituitary-adrenal (HPA) or 'hormonal' stress axis | Liu et al. 2001<sup>99</sup> |
| **Bcor** | BCL6 Corepressor | 0.0017 | Interacting partner for the BCL6 oncogenic repressor; associated with neuroepithelial tumors | Entrez Gene: BCOR, BCL6 corepressor; Bunting and Melnick 2013<sup>135</sup>; de Lima et al. 2020<sup>136</sup> |
| **Slitrk5** | SLIT and NTRK Like Family Member 5 | 0.0021 | Member of Slitrk family of proteins, involved in synaptogenesis, neurite outgrowth, neuronal survival; modulates Brain-Derived Neurotrophic Factor (BDNF)-dependent pathways via interaction with TrkB receptors; modulates hyperactivity behavior | Yim et al. 2013<sup>137</sup>; Song<sup>138</sup> 2015; Song et al. 2017<sup>139</sup>; Shmelkov et al. 2010<sup>140</sup>; Salesse et al. 2020<sup>141</sup> |
| **Rps28** | Ribosomal Protein S28 | 0.0022 | Component of the 40S (small) ribosomal subunit; likely involved in ribosome biogenesis | Kim et al. 2017<sup>142</sup> |
| **Socs5** | Suppressor Of Cytokine Signaling 5 | 0.0028 | Member of SOCS (also known as STAT-induced STAT inhibitor (SSI)) family of proteins; associated with cell proliferation and tumorigenesis | Entrez Gene: SOCS5, suppressor of cytokine signaling 5; Trengove and Ward 2013<sup>143</sup>; Sharma et al. 2019<sup>144</sup>; Sanchez-Mejias et al. 2019<sup>145</sup> |
| **Rps18** | Ribosomal Protein S18 | 0.0039 | Component of the 40S (small) ribosomal subunit; involved in initiation of translation | Entrez Gene: RPS18, ribosomal protein S18 |
| **Slc25a23** | Solute Carrier Family 25 Member 23 | 0.0052 | Calcium-dependent mitochondrial solute carrier | Bassi et al. 2005<sup>146</sup> |
| **Dnal4** | Dynein Axonemal Light Chain 4 | 0.0061 | Functions in outer dynein arms complex - a molecular motor that produces force to move cilia; associated with Congenital mirror movement disorder | Entrez Gene: DNAL4 dynein, axonemal, light chain 4; Ahmed et al. 2014<sup>147</sup> |
| **UBALD1** | UBA Like Domain Containing 1 | 0.0111 | Currently unknown |  |
| **Hmgb3** | High Mobility Group Box 3 | 0.0120 | Member of protein family with one or more high mobility group DNA-binding motifs; important role in maintaining stem cell populations | Entrez Gene: HMGB3, high mobility group box 3; Nemeth et al. 2006<sup>148</sup> |
| **Rps20** | Ribosomal Protein S20 | 0.0169 | Component of the 40S (small) ribosomal subunit; role in cell proliferation and tumor development | Krishnan et al. 2018<sup>149</sup>; Goudarzi and Lindstrom 2016<sup>150</sup> |
| **Tecpr2** | Tectonin Beta-Propeller Repeat-Containing 2 | 0.0213 | Role in autophagy-related neurodegenerative disorder spastic paraplegia type 49 (SPG49), characterized by intellectual disability, decreased pain sensitivity, autonomic-sensory neuropathy, chronic respiratory disease | Oz-Levi et al. 2012<sup>151</sup>; Tamim-Yecheskel et al. 2020<sup>152</sup> |
| **Unknown** | NA | 0.0223 | NA | NA |
| **Nipsnap3b** | Nipsnap Homolog 3B | 0.0280 | Member of protein family with possible role in vesicular trafficking | Buechler et al. 2004<sup>153</sup> |
| **Nsg2** | Neuronal Vesicle Trafficking Associated 2 | 0.0390 | Neuron-specific protein involved in modulation of excitatory neurotransmission | Chander et al. 2019<sup>154</sup> |

**Table 3.** Differentially Expressed Genes Between Stress High and Control Animals.

| Gene Symbol | Gene Name | Adjusted P-value | Related Function | References |
| --- | --- | --- | --- | --- |
| **Cartpt** | Cocaine and Amphetamine Regulated Transcript Prepropeptide | 0.0000 | Role in sensory processing, reinforcement, reward, anxiety, depression; missense mutation associated with increased anxiety in humans; hippocampal expression is modulated by stress | Hunter et al. 2007<sup>54</sup>; Subhedar et al. 2014<sup>55</sup>; Miraglia del Giudice et al. 2006<sup>57</sup>; for review see Stanek 2006<sup>56</sup> |
| ***Prdm8*** | PR/SET Domain 8 | 0.0000 | Member of conserved family of histone methyltransferases that negatively regulate transcription; involved in neural circuit assembly and neocortical development | Ross et al. 2012<sup>155</sup>; Inoue et al. 2014<sup>156</sup> |
| ***RT1-A3*** | RT1 class I, locus A3 (also MHC class I protein) | 0.0003 | Role in immune system antigen presentation; expression modulated in hippocampus after chronic immobilization stress | VanGuilder Starkey et al. 2012<sup>157</sup>; Li et al. 2013<sup>158</sup> |
| ***Hbs1l*** | Hsp70 Subfamily B Suppressor 1-Like Protein | 0.0101 | Member of GTP-binding elongation factor family; associated with regulation of fetal hemoglobin; no known CNS-specific function | Entrez Gene: HBS1L, HBS1 like translational GTPase; Stadhouders et al. 2014<sup>159</sup> |
| ***Pdyn*** | Prodynorphin | 0.0403 | Encodes a preprotein that yields several active opium-like peptides which are ligands for the kappa-type opioid receptor; ligand functions are associated to stress, anxiety, depression, pain | Entrez Gene: PDYN, prodynorphin; Baird et al. 2021<sup>160</sup>; Bodnar 2013<sup>161</sup>; Szklarczyk et al. 2012<sup>163</sup>; Johnson et al. 2021<sup>163</sup> |

## Discussion

In this study, we identified hippocampal and amygdala transcriptional signatures that correspond with sustained changes to behavior after a single, severe stressor. One week after acute stress, rats displayed substantial variability in avoidance and startle, with a subset of animals exhibiting high anxiety-like behavior. Select sets of genes corresponded with divergent anxiety-like phenotypes. Stress vulnerability strongly corresponded with upregulation of *Cartpt*, a corticosterone-regulated gene, in the hippocampal DG. In contrast, the DG of stress resilient animals displayed downregulation of genes involved in synaptic transmission and CRH signaling when compared to control or stress vulnerable animals. Transcriptional profiles of the basolateral amygdala were distinct from those of the hippocampus, with insulin/hormone signaling and anxiety-related genes largely upregulated in vulnerable animals compared to resilient animals. Altogether, these findings provide a new perspective on hippocampal and amygdala molecular regulation of divergent behavioral outcomes after trauma.

First, to measure the degree of physiological responses to stress, we measured serum corticosterone and weight loss. Rats exhibited high serum corticosterone throughout stress, and stress-exposed rats lost significantly more weight than control animals, indicating that our stress paradigm elicited a potent stress response. After 1 week of recovery from stress, rats underwent a multivariate behavioral assessment. We have shown that this paradigm reveals a spectrum of avoidance, startle, and fear learning behaviors in individual rats^19^, which was replicated in this cohort. We tested whether physiological responses to acute stress could predict future behavioral outcomes. Interestingly, corticosterone at the end, but not the beginning, of the 3-hour stressor positively correlated with anxiety-like behavior scores one week later. Corticosterone was not a perfect predictor, as only 30% of the variance of behavior scores could be explained by corticosterone. Nonetheless, this may suggest that sustained release of corticosterone throughout a traumatic event contributes to structural and functional changes leading to enhanced or persistent anxiety-like behavior^38–40^. This finding contrasts with several studies suggesting that corticosterone measured soon after trauma is negatively correlated with PTSD outcomes^41,42^.

We identified transcriptional signatures that correspond with behavioral outcomes after stress. Principal component analysis indicated that individual samples did not clearly cluster by condition, suggesting that global gene expression of both the dentate gyrus and amygdala does not correlate with persistent anxiety-like behavior after acute stress. Rather, differences in behavioral outcomes may be attributable to changes among select sets of genes. In the amygdala, comparisons between control and stress-vulnerable, and control and stress-resilient groups returned few genes (10 for both) with a large overlap between the 2 sets of genes - indicating a largely homogeneous response. Between vulnerable and resilient animals, we found 22 DEGs that enriched for biological processes pertaining to the response to insulin or hormone stimulus. Interestingly, impaired insulin signaling in the amygdala has been shown to affect synaptic function and to increase anxiety-like behavior in mice. Specifically, deficient insulin signaling in the central amygdala via knockout of both insulin receptors (insulin receptor (IR) and insulin-like growth factor 1 receptor (IGF1R)) has been shown to alter synaptic function, impair cognition, and increase anxiety-like behavior^43^. Similarly, associations between insulin resistance and mood disorders (i.e., anxiety) have been reported in humans^44^. In our study, 3 of the top 2 biological process category genes (*Inhbb, Rab31, Ncoa3)*, which also have functions directly associated with modulation of anxiety-like behaviors in mice, were upregulated in vulnerable animals compared to resilient animals.

Interestingly, IGF-1 (which also signals through IR and IGF1R) is involved in mood regulation^45,46^. IGF-1 administration in rats has been shown to mimic the effects of early-life environmental enrichment, increase glucocorticoid receptor (GR) levels in the hippocampus and reduce anxiety-like behavior in adults^47^. Thus, IGF-1 signaling may mediate some of the long-term effects of early-life experiences on stress resilience/vulnerability later in life. Similarly, intracerebroventricular injection of IGF-1 in mice decreased anxiety in the EPM and open-field after predator exposure – an effect mediated in part via stimulation of FKBP5 expression (an important modulator of GR signaling and neuroendocrine stress responses^48,49^. It is interesting to consider whether similar mechanisms might underlie adaptations in the amygdala following severe acute stress exposure.

Transcriptional profiles of the hippocampal DG were distinct from those of the amygdala in vulnerable and resilient animals, with essentially no overlap of differentially expressed genes. In the DG, our analyses returned 40 genes for the resilient vs vulnerable comparison, 24 genes for the control vs resilient comparison, and 5 genes for the control vs vulnerable comparison. Hence, very few genes distinguish control animals from vulnerable animals. Instead, greater numbers of genes separate resilient animals from both control and vulnerable animals, which may suggest that an active transcriptional response is linked to resilience.

It is worth noting that, in the DG, various DEGs in the resilient vs vulnerable comparison (shown in Table 1) have roles related to dendritic spine structure and function (*Rnf112, Snap25*), as well as neurite outgrowth and arborization (*Tnk2, Dpysl5, Rtn3, Rtn4*). Dendritic remodeling (including dendritic extension, retraction, and arborization, as well as changes in spine number, shape, size, and stability) in brain regions responsive to stress has been extensively studied and is a crucial mechanism underlying stress-induced changes in synaptic function, plasticity, and connectivity. These in turn are thought to contribute to the expression of adaptive or maladaptive stress-related behaviors including anxiety (for review see^20,50^). We also found differentially expressed genes involved in anxiety, mood, emotion (*Cartpt, Snap25, Rxfp1, Nr1d1, Sept5*), and neuropsychiatric disorders (YWHA-family). In many cases, expression patterns of these genes were up- or downregulated specifically in the stress-resilient group, suggesting that resilience is not a passive lack of response, but an active means of protection from the effects of stress.

Additionally, the overlapping set of genes between control vs resilient and vulnerable vs resilient comparisons in the DG (‘resilience signature’) indicated that *Rnf112* and *Tbx19* are significantly downregulated in resilient animals. Knockout of *Rnf112* has been shown to decrease excitatory synaptic transmission through modulation of spine density and excitatory synapse function in mice^51^. Importantly, decreased excitatory synaptic transmission in the DG (via NR1 NMDA receptor subunit deletion) reduces anxiety in mice and may be a key locus mediating the anxiolytic effects of NMDA receptor antagonists^52^. *Tbx19* is involved in the corticotropin-releasing hormone (CRH) signaling pathway - a central part of the hypothalamic-pituitary-adrenal (HPA) axis. While the function of *Tbx19* specifically in the hippocampus is currently unknown, hippocampal CRH signaling does modulate synaptic function and plasticity - a bidirectional regulation which is dependent on stress duration and severity^53^.

Finally, between the resilient vs vulnerable and control vs vulnerable comparisons, we found a single overlapping gene – *Cartpt*, expression of which significantly correlated with composite behavior scores. *Cartpt* encodes a multifaceted neuropeptide that is highly expressed in limbic regions. Stress modulates *Cartpt* expression, and *Cartpt* has well-established roles in reinforcement, reward, sensory processing, anxiety, and depression (among others)^54^. Importantly, several studies have shown that *Cartpt* levels are increased in animal models of anxiety^55,56^, and a missense mutation in the *Cartpt* gene has been associated with increased anxiety in humans^57^. Upregulated *Cartpt* in the DG of animals displaying high anxiety scores might thus serve as an indicator of susceptibility after an acute, severe stressor.

One limitation of our study is that, because our experimental design is cross-sectional in behavior and transcription measures, our results do not indicate when changes to gene expression first occur. Thus, a critical future direction for this work will be to determine the time course of transcriptional regulation such that key genes may be targeted to prevent or relieve anxiety-like behavior after stress. In addition, our results are specific to males, and future experiments should expand this work to include females.

In conclusion, we demonstrate that divergent behavioral outcomes after a single, severe stress are associated with regionally distinct transcriptional profiles in the amygdala and the hippocampus. Key genes corresponding with resilience and vulnerability in the amygdala are associated with insulin and hormonal signaling. In the hippocampus, *Cartpt* was strongly correlated with vulnerability to anxiety-like behavior; however, transcriptional changes were primarily associated with stress resilience, suggesting that an active molecular response in the hippocampus facilitates protection from the long-term consequences of severe stress. These results provide novel insight into the mechanisms that bring about individual variability in the behavioral responses to stress and provide new targets for the advancement of therapies for stress-induced neuropsychiatric disorders.

## Acknowledgments

We thank Yurika Kazama, Rhea Misra, Claire Toth, and Kelsey Hu for assistance with animal work and data collection. Genomic library preparation and sequencing was performed by the Vincent J. Coates Genomics Sequencing Laboratory at UC Berkeley.

## Data and materials availability

All data associated with this study are available upon request.

## Supplementary Materials

Fig. S1: PCA of normalized gene expression values for the amygdala and hippocampus dentate gyrus (DG) samples.

Fig. S2: All behavior data.

Fig. S3: Behavioral correlations and clustering.

Fig. S4: Correlations between corticosterone, weight loss, and behavior after stress.

Fig. S5: *Rarres2* gene expression correlations to Overall Composite, Avoidance Composite and Startle Composite Score.

Fig. S6: *ATN1* and *CTTN* gene expression differences between control, stress-low and stress-high groups.

Fig. S7: *Cartpt* gene expression correlations to Overall Composite, Avoidance Composite and Startle Composite Score

## References

1. Yehuda, R. & LeDoux, J. Response Variation following Trauma: A Translational Neuroscience Approach to Understanding PTSD. Neuron 56, 19–32 (2007).

2. Kessler, R. C. Posttraumatic Stress Disorder in the National Comorbidity Survey. Arch. Gen. Psychiatry 52, 1048 (1995).

3. Jin, Y., Xu, J., Liu, H. & Liu, D. Posttraumatic stress disorder and posttraumatic growth among adult survivors of Wenchuan earthquake after 1 year: prevalence and correlates. Arch. Psychiatr. Nurs. 28, 67–73 (2014).

4. Roozendaal, B., McEwen, B. S. & Chattarji, S. Stress, memory and the amygdala. Nat. Rev. Neurosci. 10, 423–433 (2009).

5. McEwen, B. S. Stress and Hippocampal Plasticity. Annu. Rev. Neurosci. 22, 105–122 (1999).

6. Davis, M. The Role of the Amygdala in Fear and Anxiety. Annu. Rev. Neurosci. 15, 353–375 (1992).

7. Phillips, R. G. & LeDoux, J. E. Differential contribution of amygdala and hippocampus to cued and contextual fear conditioning. Behav. Neurosci. 106, 274–285 (1992).

8. Shi, H.-J., Wang, S., Wang, X.-P., Zhang, R.-X. & Zhu, L.-J. Hippocampus: Molecular, Cellular, and Circuit Features in Anxiety. Neurosci. Bull. (023) doi:10.1007/s12264-023-01020-1.

9. Jankord, R. & Herman, J. P. Limbic regulation of hypothalamo-pituitary-adrenocortical function during acute and chronic stress. Ann. N. Y. Acad. Sci. 1148, 64–73 (2008).

10. McEwen, B. S., Nasca, C. & Gray, J. D. Stress Effects on Neuronal Structure: Hippocampus, Amygdala and Prefrontal Cortex. Neuropsychopharmacology 41, 3–23 (2016).

11. von Ziegler, L. M. et al. Multiomic profiling of the acute stress response in the mouse hippocampus. Nat. Commun. 13, 1824 (2022).

12. Lori, A. et al. Dynamic Patterns of Threat-Associated Gene Expression in the Amygdala and Blood. Front. Psychiatry 9, (2019).

13. Sillivan, S. E., Jones, M. E., Jamieson, S., Rumbaugh, G. & Miller, C. A. Bioinformatic analysis of long-lasting transcriptional and translational changes in the basolateral amygdala following acute stress. PLOS ONE 14, e0209846 (2019).

14. Shen, M., Song, Z. & Wang, J.-H. microRNA and mRNA profiles in the amygdala are associated with stress-induced depression and resilience in juvenile mice. Psychopharmacology (Berl.) 236, 2119–2142 (2019).

15. Ponomarev, I., Rau, V., Eger, E. I., Harris, R. A. & Fanselow, M. S. Amygdala transcriptome and cellular mechanisms underlying stress-enhanced fear learning in a rat model of posttraumatic stress disorder. Neuropsychopharmacol. Off. Publ. Am. Coll. Neuropsychopharmacol. 35, 1402–1411 (2010).

16. Fa, M. et al. Stress modulation of hippocampal activity – Spotlight on the dentate gyrus. Neurobiol. Learn. Mem. 112, 53–60 (2014).

17. Datson, N. A. et al. Previous History of Chronic Stress Changes the Transcriptional Response to Glucocorticoid Challenge in the Dentate Gyrus Region of the Male Rat Hippocampus. Endocrinology 154, 3261–3272 (2013).

18. Kirby, E. D. et al. Acute stress enhances adult rat hippocampal neurogenesis and activation of newborn neurons via secreted astrocytic FGF2. eLife 2, 1–23 (2013).

19. Long, K. L. P. et al. Regional gray matter oligodendrocyte- and myelin-related measures are associated with differential susceptibility to stress-induced behavior in rats and humans. Transl. Psychiatry 11, 1–15 (2021).

20. Vyas, A., Mitra, R., Rao, B. S. S. & Chattarji, S. Chronic Stress Induces Contrasting Patterns of Dendritic Remodeling in Hippocampal and Amygdaloid Neurons. J. Neurosci. 22, 6810–6818 (2002).

21. Mitra, R., Jadhav, S., McEwen, B. S., Vyas, A. & Chattarji, S. Stress duration modulates the spatiotemporal patterns of spine formation in the basolateral amygdala. Proc. Natl. Acad. Sci. U. S. A. 102, 9371–9376 (2005).

22. Sillivan, S. E. et al. Susceptibility and Resilience to Posttraumatic Stress Disorder–like Behaviors in Inbred Mice. Biol. Psychiatry 82, 924–933 (2017).

23. Cohen, H. & Zohar, J. An Animal Model of Posttraumatic Stress Disorder: The Use of Cut-Off Behavioral Criteria. Ann. N. Y. Acad. Sci. 1032, 167–178 (2004).

24. Dobin, A. et al. STAR: ultrafast universal RNA-seq aligner. Bioinformatics 29, 15–21 (2012).

25. Li, B. & Dewey, C. N. RSEM: accurate transcript quantification from RNA-Seq data with or without a reference genome. BMC Bioinformatics 12, 323 (2011).

26. Leng, N. et al. EBSeq: an empirical Bayes hierarchical model for inference in RNA-seq experiments. Bioinformatics 29, 1035–1043 (2013).

27. Mootha, V. K. et al. PGC-1α-responsive genes involved in oxidative phosphorylation are coordinately downregulated in human diabetes. Nat. Genet. 34, 267–273 (2003).

28. Subramanian, A. et al. Gene set enrichment analysis: a knowledge-based approach for interpreting genome-wide expression profiles. Proc. Natl. Acad. Sci. U. S. A. 102, 15545–15550 (2005).

29. Muroy, S. E., Long, K. L. P., Kaufer, D. & Kirby, E. D. Moderate Stress-Induced Social Bonding and Oxytocin Signaling are Disrupted by Predator Odor in Male Rats. Neuropsychopharmacology 41, 2160–2170 (2016).

30. Breton, J. M. et al. Juvenile exposure to acute traumatic stress leads to long-lasting alterations in grey matter myelination in adult female but not male rats. Neurobiol. Stress 14, 100319 (2021).

31. Ageta, H. et al. Activin in the Brain Modulates Anxiety-Related Behavior and Adult Neurogenesis. PLoS ONE 3, e1869 (2008).

32. Zheng, F. et al. Activin tunes GABAergic neurotransmission and modulates anxiety-like behavior. Mol. Psychiatry 14, 332–346 (2009).

33. Link, A. S., Zheng, F. & Alzheimer, C. Activin Signaling in the Pathogenesis and Therapy of Neuropsychiatric Diseases. Front. Mol. Neurosci. 9, 32 (2016).

34. Rouillard, A. D. et al. The harmonizome: a collection of processed datasets gathered to serve and mine knowledge about genes and proteins. Database 2016, baw100 (2016).

35. Sun, Z. & Xu, Y. Nuclear Receptor Coactivators (NCOAs) and Corepressors (NCORs) in the Brain. Endocrinology 161, bqaa083 (2020).

36. Stashi, E., Wang, L., Mani, S. K., York, B. & O’Malley, B. W. Research resource: loss of the steroid receptor coactivators confers neurobehavioral consequences. Mol. Endocrinol. Baltim. Md 27, 1776–1787 (2013).

37. York, B. et al. Research resource: tissue- and pathway-specific metabolomic profiles of the steroid receptor coactivator (SRC) family. Mol. Endocrinol. Baltim. Md 27, 366–380 (2013).

38. McEwen, B. S. Glucocorticoids, depression, and mood disorders: structural remodeling in the brain. Metabolism 54, 20–23 (2005).

39. Chetty, S. et al. Stress and glucocorticoids promote oligodendrogenesis in the adult hippocampus. Mol. Psychiatry 19, 1275–1283 (2014).

40. Chen, Y., Dube, C. M., Rice, C. J. & Baram, T. Z. Rapid Loss of Dendritic Spines after Stress Involves Derangement of Spine Dynamics by Corticotropin-Releasing Hormone. J. Neurosci. 28, 2903–2911 (2008).

41. Cohen, H. et al. Blunted HPA Axis Response to Stress Influences Susceptibility to Posttraumatic Stress Response in Rats. Biol. Psychiatry 59, 1208–1218 (2006).

42. Yehuda, R. Current status of cortisol findings in post-traumatic stress disorder. Psychiatr. Clin. North Am. 25, 341–68, vii (2002).

43. Soto, M., Cai, W., Konishi, M. & Kahn, C. R. Insulin signaling in the hippocampus and amygdala regulates metabolism and neurobehavior. Proc. Natl. Acad. Sci. 116, 6379–6384 (2019).

44. Li, C. et al. Diabetes and anxiety in US adults: findings from the 2006 Behavioral Risk Factor Surveillance System. Diabet. Med. J. Br. Diabet. Assoc. 25, 878–881 (2008).

45. Hoshaw, B. A. et al. Antidepressant-like behavioral effects of IGF-I produced by enhanced serotonin transmission. Eur. J. Pharmacol. 594, 109–116 (2008).

46. Bot, M., Milaneschi, Y., Penninx, B. W. J. H. & Drent, M. L. Plasma insulin-like growth factor I levels are higher in depressive and anxiety disorders, but lower in antidepressant medication users. Psychoneuroendocrinology 68, 148–155 (2016).

47. Baldini, S. et al. Enriched Early Life Experiences Reduce Adult Anxiety-Like Behavior in Rats: A Role for Insulin-Like Growth Factor 1. J. Neurosci. 33, 11715–11723 (2013).

48. Zannas, A. S., Wiechmann, T., Gassen, N. C. & Binder, E. B. Gene–Stress–Epigenetic Regulation of FKBP5: Clinical and Translational Implications. Neuropsychopharmacology 41, 261–274 (2016).

49. Santi, A., Bot, M., Aleman, A., Penninx, B. W. J. H. & Aleman, I. T. Circulating insulin-like growth factor I modulates mood and is a biomarker of vulnerability to stress: from mouse to man. Transl. Psychiatry 8, 1–11 (2018).

50. Leuner, B. & Shors, T. J. Stress, anxiety, and dendritic spines: What are the connections? Neuroscience 251, 108–119 (2013).

51. Lomash, R. M., Gu, X., Youle, R. J., Lu, W. & Roche, K. W. Neurolastin, a Dynamin Family GTPase, Regulates Excitatory Synapses and Spine Density. Cell Rep. 12, 743–751 (2015).

52. Barkus, C. et al. Hippocampal NMDA receptors and anxiety: At the interface between cognition and emotion. Eur. J. Pharmacol. 626, 49–56 (2010).

53. Chen, Y., Andres, A. L., Frotscher, M. & Baram, T. Z. Tuning synaptic transmission in the hippocampus by stress: the CRH system. Front. Cell. Neurosci. 6, 13 (2012).

54. Hunter, R. G. et al. Regulation of CART mRNA by Stress and Corticosteroids in the Hippocampus and Amygdala. Brain Res. 1152, 234–240 (2007).

55. Subhedar, N. K., Nakhate, K. T., Upadhya, M. A. & Kokare, D. M. CART in the brain of vertebrates: Circuits, functions and evolution. Peptides 54, 108–130 (2014).

56. Stanek, L. M. Cocaine- and amphetamine related transcript (CART) and anxiety. Peptides 27, 2005–2011 (2006).

57. Miraglia del Giudice, E. et al. Adolescents carrying a missense mutation in the CART gene exhibit increased anxiety and depression. Depress. Anxiety 23, 90–92 (2006).

58. Alicea, D., Perez, M., Maldonado, C., Dominicci-Cotto, C. & Marie, B. Cortactin Is a Regulator of Activity-Dependent Synaptic Plasticity Controlled by Wingless. J. Neurosci. 37, 2203–2215 (2017).

59. Cornelius, J., Rottner, K., Korte, M. & Michaelsen-Preusse, K. Cortactin Contributes to Activity-Dependent Modulation of Spine Actin Dynamics and Spatial Memory Formation. Cells 10, 1835 (2021).

60. Cornell, B. & Toyo-oka, K. 14-3-3 Proteins in Brain Development: Neurogenesis, Neuronal Migration and Neuromorphogenesis. Front. Mol. Neurosci. 0, (2017).

61. Berg, D., Holzmann, C. & Riess, O. 14-3-3 proteins in the nervous system. Nat. Rev. Neurosci. 4, 752–762 (2003).

62. Foote, M. & Zhou, Y. 14-3-3 proteins in neurological disorders. Int. J. Biochem. Mol. Biol. 3, 152–164 (2012).

63. Shi, Q., Ge, Y., He, W., Hu, X. & Yan, R. RTN1 and RTN3 protein are differentially associated with senile plaques in Alzheimer’s brains. Sci. Rep. 7, 6145 (2017).

64. Alhajlah, S., Thompson, A. M. & Ahmed, Z. Overexpression of Reticulon 3 enhances CNS axon regeneration and functional recovery after injury. bioRxiv 2021.05.24.445420 (2021) doi:10.1101/2021.05.24.445420.

65. Oertle, T. & Schwab, M. E. Nogo and its paRTNers. Trends Cell Biol. 13, 187–194 (2003).

66. Wood, J. D. et al. Atrophin-1, the Dentato-Rubral and Pallido-Luysian Atrophy Gene Product, Interacts with Eto/Mtg8 in the Nuclear Matrix and Represses Transcription. J. Cell Biol. 150, 939–948 (2000).

67. Shen, Y. & Peterson, A. S. Atrophins’ emerging roles in development and neurodegenerative disease. Cell. Mol. Life Sci. 66, 437 (2008).

68. Suzuki, G. et al. Sept5 deficiency exerts pleiotropic influence on affective behaviors and cognitive functions in mice. Hum. Mol. Genet. 18, 1652–1660 (2009).

69. Gao, Y.-Y. et al. Recombinant milk fat globule-EGF factor-8 reduces apoptosis via integrin β3/FAK/PI3K/AKT signaling pathway in rats after traumatic brain injury. Cell Death Dis. 9, 1–17 (2018).

70. Cheyuo, C., Aziz, M. & Wang, P. Neurogenesis in Neurodegenerative Diseases: Role of MFG-E8. Front. Neurosci. 13, (2019).

71. Cahoy, J. D. et al. A Transcriptome Database for Astrocytes, Neurons, and Oligodendrocytes: A New Resource for Understanding Brain Development and Function. J. Neurosci. 28, 264–278 (2008).

72. Hayne, C. K., Schmidt, C. A., Haque, M. I., Matera, A. G. & Stanley, R. E. Reconstitution of the human tRNA splicing endonuclease complex: insight into the regulation of pre-tRNA cleavage. Nucleic Acids Res. 48, 7609–7622 (2020).

73. Namavar, Y. et al. Clinical, neuroradiological and genetic findings in pontocerebellar hypoplasia. Brain J. Neurol. 134, 143–156 (2011).

74. Matsuda, Y. et al. Chromosome Mapping of Human (ZNF179), Mouse, and Rat Genes for Brain Finger Protein (bfp), a Member of the RING Finger Family. Genomics 33, 325–327 (1996).

75. Zhou, Y., Gunput, R.-A. F., Adolfs, Y. & Pasterkamp, R. J. MICALs in control of the cytoskeleton, exocytosis, and cell death. Cell. Mol. Life Sci. CMLS 68, 4033–4044 (2011).

76. Jiang, P. et al. A Systems Approach Identifies Networks and Genes Linking Sleep and Stress: Implications for Neuropsychiatric Disorders. Cell Rep. 11, 835–848 (2015).

77. Bagchi, S. et al. Probable role for major facilitator superfamily domain containing 6 (MFSD6) in the brain during variable energy consumption. Int. J. Neurosci. 130, 476–489 (2020).

78. McCulloch, K. A., Qi, Y. B., Takayanagi-Kiya, S., Jin, Y. & Cherra, S. J. Novel Mutations in Synaptic Transmission Genes Suppress Neuronal Hyperexcitation in Caenorhabditis elegans. G3 GenesGenomesGenetics 7, 2055–2063 (2017).

79. Choi, H. et al. Acetylation changes tau interactome to degrade tau in Alzheimer’s disease animal and organoid models. Aging Cell 19, e13081 (2020).

80. Gootjes, J., Schmohl, F., Waterham, H. R. & Wanders, R. J. A. Novel mutations in the PEX12 gene of patients with a peroxisome biogenesis disorder. Eur. J. Hum. Genet. 12, 115–120 (2004).

81. Steinberg, S. J. et al. Peroxisome biogenesis disorders. Biochim. Biophys. Acta BBA - Mol. Cell Res. 1763, 1733–1748 (2006).

82. Steinberg, S. J., Raymond, G. V., Braverman, N. E. & Moser, A. B. Zellweger Spectrum Disorder. in GeneReviews® (eds. Adam, M. P. et al.) (University of Washington, Seattle, 1993).

83. Steinberg, S. et al. The PEX Gene Screen: molecular diagnosis of peroxisome biogenesis disorders in the Zellweger syndrome spectrum. Mol. Genet. Metab. 83, 252–263 (2004).

84. Ureña, J. M. et al. Expression, synaptic localization, and developmental regulation of Ack1/Pyk1, a cytoplasmic tyrosine kinase highly expressed in the developing and adult brain. J. Comp. Neurol. 490, 119–132 (2005).

85. La Torre, A., del Rio, J. A., Soriano, E. & Ureña, J. M. Expression pattern of ACK1 tyrosine kinase during brain development in the mouse. Gene Expr. Patterns 6, 886–892 (2006).

86. La Torre, A. et al. A role for the tyrosine kinase ACK1 in neurotrophin signaling and neuronal extension and branching. Cell Death Dis. 4, e602 (2013).

87. Hitomi, Y. et al. Mutations in TNK2 in severe autosomal recessive infantile onset epilepsy. Ann. Neurol. 74, 496–501 (2013).

88. Chiricosta, L., Gugliandolo, A., Bramanti, P. & Mazzon, E. Could the Heat Shock Proteins 70 Family Members Exacerbate the Immune Response in Multiple Sclerosis? An in Silico Study. Genes 11, 615 (2020).

89. Zhao, C. & Gammie, S. C. The circadian gene Nr1d1 in the mouse nucleus accumbens modulates sociability and anxiety-related behaviour. Eur. J. Neurosci. 48, 1924–1943 (2018).

90. Smagin, D. A. et al. Dysfunction in Ribosomal Gene Expression in the Hypothalamus and Hippocampus following Chronic Social Defeat Stress in Male Mice as Revealed by RNA-Seq. Neural Plast. 2016, e3289187 (2015).

91. Lee, J. H. et al. Altered relaxin family receptors RXFP1 and RXFP3 in the neocortex of depressed Alzheimer’s disease patients. Psychopharmacology (Berl.) 233, 591–598 (2016).

92. Kataoka, M. et al. A Single Amino Acid Mutation in SNAP-25 Induces Anxiety-Related Behavior in Mouse. PLOS ONE 6, e25158 (2011).

93. Irfan, M. et al. SNAP-25 isoforms differentially regulate synaptic transmission and long-term synaptic plasticity at central synapses. Sci. Rep. 9, 6403 (2019).

94. Karthikeyan, S. et al. Crystal structure of human riboflavin kinase reveals a beta barrel fold and a novel active site arch. Struct. Lond. Engl. 1993 11, 265–273 (2003).

95. Kukharsky, M. S. et al. Long non-coding RNA Neat1 regulates adaptive behavioural response to stress in mice. Transl. Psychiatry 10, 1–19 (2020).

96. Johnson, J. O. et al. Exome sequencing reveals riboflavin transporter mutations as a cause of motor neuron disease. Brain J. Neurol. 135, 2875–2882 (2012).

97. Hirano, A., Braas, D., Fu, Y.-H. & Ptáček, L. J. FAD Regulates CRYPTOCHROME Protein Stability and Circadian Clock in Mice. Cell Rep. 19, 255–266 (2017).

98. Shan, A. et al. Polypeptide N-acetylgalactosaminyltransferase 18 non-catalytically regulates the ER homeostasis and O-glycosylation. Biochim. Biophys. Acta Gen. Subj. 1863, 870–882 (2019).

99. Liu, J. et al. Tbx19, a tissue-selective regulator of POMC gene expression. Proc. Natl. Acad. Sci. U. S. A. 98, 8674–8679 (2001).

100. Lissandron, V., Podini, P., Pizzo, P. & Pozzan, T. Unique characteristics of Ca2+ homeostasis of the trans-Golgi compartment. Proc. Natl. Acad. Sci. 107, 9198–9203 (2010).

101. Berg, D., Holzmann, C. & Riess, O. 14-3-3 proteins in the nervous system. Nat. Rev. Neurosci. 4, 752–762 (2003).

102. Foote, M. & Zhou, Y. 14-3-3 proteins in neurological disorders. Int. J. Biochem. Mol. Biol. 3, 152–164 (2012).

103. Sillivan, S. E., Jones, M. E., Jamieson, S., Rumbaugh, G. & Miller, C. A. Bioinformatic analysis of long-lasting transcriptional and translational changes in the basolateral amygdala following acute stress. PLoS ONE 14, e0209846 (2019).

104. Thiffault, I. et al. Pathogenic variants in KPTN gene identified by clinical whole-genome sequencing. Mol. Case Stud. 6, a003970 (2020).

105. Pajusalu, S., Reimand, T. & Õdunap, K. Novel homozygous mutation in KPTN gene causing a familial intellectual disability-macrocephaly syndrome. Am. J. Med. Genet. A. 167A, 1913–1915 (2015).

106. Chen, Y. et al. Wdr47 Controls Neuronal Polarization through the Camsap Family Microtubule Minus-End-Binding Proteins. Cell Rep. 31, 107526 (2020).

107. Kannan, M. et al. WD40-repeat 47, a microtubule-associated protein, is essential for brain development and autophagy. Proc. Natl. Acad. Sci. 114, E9308–E9317 (2017).

108. GrandPré, T., Nakamura, F., Vartanian, T. & Strittmatter, S. M. Identification of the Nogo inhibitor of axon regeneration as a Reticulon protein. Nature 403, 439–444 (2000).

109. Petrinovic, M. M. et al. Neuronal Nogo-A regulates neurite fasciculation, branching and extension in the developing nervous system. Development 137, 2539–2550 (2010).

110. Wälchli, T. et al. Nogo-A is a negative regulator of CNS angiogenesis. Proc. Natl. Acad. Sci. 110, E1943–E1952 (2013).

111. Acevedo, L. et al. A new role for Nogo as a regulator of vascular remodeling. Nat. Med. 10, 382–388 (2004).

112. Jia, S. et al. Nogo-C regulates cardiomyocyte apoptosis during mouse myocardial infarction. Cell Death Dis. 7, e2432–e2432 (2016).

113. Liu, K. X. et al. Neuron-specific antioxidant OXR1 extends survival of a mouse model of amyotrophic lateral sclerosis. Brain J. Neurol. 138, 1167–1181 (2015).

114. Volkert, M. R. & Crowley, D. J. Preventing Neurodegeneration by Controlling Oxidative Stress: The Role of OXR1. Front. Neurosci. 14, 1318 (2020).

115. Jeanne, M. et al. Missense variants in DPYSL5 cause a neurodevelopmental disorder with corpus callosum agenesis and cerebellar abnormalities. Am. J. Hum. Genet. 108, 951–961 (2021).

116. Quach, T. T., Honnorat, J., Kolattukudy, P. E., Khanna, R. & Duchemin, A. M. CRMPs: critical molecules for neurite morphogenesis and neuropsychiatric diseases. Mol. Psychiatry 20, 1037–1045 (2015).

117. Lin, Y.-F., Chen, K. C., Yang, Y. K. & Hsiao, Y.-H. Collapsin response mediator protein 5 (CRMP5) modulates susceptibility to chronic social defeat stress in mice. Mol. Neurobiol. 58, 3175–3186 (2021).

118. Rebhan, M., Chalifa-Caspi, V., Prilusky, J. & Lancet, D. GeneCards: integrating information about genes, proteins and diseases. Trends Genet. 13, 163 (1997).

119. Son, H. G. et al. Prefoldin 6 mediates longevity response from heat shock factor 1 to FOXO in C. elegans. Genes Dev. 32, 1562–1575 (2018).

120. Kelly, S. D., Harrell, C. S. & Neigh, G. N. Chronic Stress Modulates Regional Cerebral Glucose Transporter Expression in an Age-Specific and Sexually-Dimorphic Manner. Physiol. Behav. 126, 39–49 (2014).

121. Bieging-Rolett, K. T. et al. Zmat3 Is a Key Splicing Regulator in the p53 Tumor Suppression Program. Mol. Cell 80, 452-469.e9 (2020).

122. Vilborg, A., Bersani, C., Wilhelm, M. T. & Wiman, K. G. The p53 target Wig-1: a regulator of mRNA stability and stem cell fate? Cell Death Differ. 18, 1434–1440 (2011).

123. Gillardon, F., Spranger, M., Tiesler, C. & Hossmann, K. A. Expression of cell death-associated phospho-c-Jun and p53-activated gene 608 in hippocampal CA1 neurons following global ischemia. Brain Res. Mol. Brain Res. 73, 138–143 (1999).

124. Asanuma, M. et al. Suppression of p53-activated gene, PAG608, attenuates methamphetamine-induced neurotoxicity. Neurosci. Lett. 414, 263–267 (2007).

125. Morimoto, N. et al. Induction of parkinsonism-related proteins in the spinal motor neurons of transgenic mouse carrying a mutant SOD1 gene. J. Neurosci. Res. 88, 1804–1811 (2010).

126. Shimizu, M. et al. Specific induction of PAG608 in cranial and spinal motor neurons of L-DOPA-treated parkinsonian rats. Neurosci. Res. 60, 355–363 (2008).

127. Ehrhardt, A., Wang, B., Leung, M. J. & Schrader, J. W. Absence of M-Ras modulates social behavior in mice. BMC Neurosci. 16, 68 (2015).

128. Engelhard, C. et al. MEGF8 is a modifier of BMP signaling in trigeminal sensory neurons. eLife 2, e01160 (2013).

129. Gunduz-Cinar, O., Hill, M. N., McEwen, B. S. & Holmes, A. Amygdala FAAH and anandamide: mediating protection and recovery from stress. Trends Pharmacol. Sci. 34, 637–644 (2013).

130. Morena, M. et al. Upregulation of Anandamide Hydrolysis in the Basolateral Complex of Amygdala Reduces Fear Memory Expression and Indices of Stress and Anxiety. J. Neurosci. Off. J. Soc. Neurosci. 39, 1275–1292 (2019).

131. Weber, H. et al. SPOCK3, a risk gene for adult ADHD and personality disorders. Eur. Arch. Psychiatry Clin. Neurosci. 264, 409–421 (2014).

132. Smagin, D. A. et al. Dysfunction in Ribosomal Gene Expression in the Hypothalamus and Hippocampus following Chronic Social Defeat Stress in Male Mice as Revealed by RNA-Seq. Neural Plast. 2016, e3289187 (2015).

133. Erro, R. Familial Paroxysmal Nonkinesigenic Dyskinesia. in GeneReviews® (eds. Adam, M. P. et al.) (University of Washington, Seattle, 1993).

134. Sun, N. et al. The PNKD gene is associated with Tourette Disorder or Tic disorder in a multiplex family. Mol. Psychiatry 23, 1487–1495 (2018).

135. Bunting, K. L. & Melnick, A. M. New effector functions and regulatory mechanisms of BCL6 in normal and malignant lymphocytes. Curr. Opin. Immunol. 25, 339–346 (2013).

136. De Lima, L. et al. Central nervous system high-grade neuroepithelial tumor with BCOR alteration (CNS HGNET-BCOR)-case-based reviews. Childs Nerv. Syst. ChNS Off. J. Int. Soc. Pediatr. Neurosurg. 36, 1589–1599 (2020).

137. Yim, Y. S. et al. Slitrks control excitatory and inhibitory synapse formation with LAR receptor protein tyrosine phosphatases. Proc. Natl. Acad. Sci. U. S. A. 110, 4057–4062 (2013).

138. Song, M. et al. Slitrk5 Mediates BDNF-Dependent TrkB Receptor Trafficking and Signaling. Dev. Cell 33, 690–702 (2015).

139. Song, M. et al. Rare Synaptogenesis-Impairing Mutations in SLITRK5 Are Associated with Obsessive Compulsive Disorder. PloS One 12, e0169994 (2017).

140. Shmelkov, S. V. et al. Slitrk5 deficiency impairs corticostriatal circuitry and leads to obsessive-compulsive-like behaviors in mice. Nat. Med. 16, 598–602, 1p following 602 (2010).

141. Salesse, C. et al. Opposite Control of Excitatory and Inhibitory Synapse Formation by Slitrk2 and Slitrk5 on Dopamine Neurons Modulates Hyperactivity Behavior. Cell Rep. 30, 2374-2386.e5 (2020).

142. Kim, H. K. et al. A transfer-RNA-derived small RNA regulates ribosome biogenesis. Nature 552, 57–62 (2017).

143. Trengove, M. C. & Ward, A. C. SOCS proteins in development and disease. Am. J. Clin. Exp. Immunol. 2, 1–29 (2013).

144. Sharma, N. D. et al. Epigenetic silencing of SOCS5 potentiates JAK-STAT signaling and progression of T-cell acute lymphoblastic leukemia. Cancer Sci. 110, 1931–1946 (2019).

145. Sanchez-Mejias, A. et al. A novel SOCS5/miR-18/miR-25 axis promotes tumorigenesis in liver cancer. Int. J. Cancer 144, 311–321 (2019).

146. Bassi, M. T. et al. Cellular expression and alternative splicing of SLC25A23, a member of the mitochondrial Ca2+-dependent solute carrier gene family. Gene 345, 173–182 (2005).

147. Ahmed, I. et al. Identification of a homozygous splice site mutation in the dynein axonemal light chain 4 gene on 22q13.1 in a large consanguineous family from Pakistan with congenital mirror movement disorder. Hum. Genet. 133, 1419–1429 (2014).

148. Nemeth, M. J., Kirby, M. R. & Bodine, D. M. Hmgb3 regulates the balance between hematopoietic stem cell self-renewal and differentiation. Proc. Natl. Acad. Sci. 103, 13783–13788 (2006).

149. Krishnan, R., Boddapati, N. & Mahalingam, S. Interplay between human nucleolar GNL1 and RPS20 is critical to modulate cell proliferation. Sci. Rep. 8, 11421 (2018).

150. Goudarzi, K. M. & Lindström, M. S. Role of ribosomal protein mutations in tumor development (Review). Int. J. Oncol. 48, 1313–1324 (2016).

151. Oz-Levi, D. et al. Mutation in TECPR2 reveals a role for autophagy in hereditary spastic paraparesis. Am. J. Hum. Genet. 91, 1065–1072 (2012).

152. Tamim-Yecheskel, B.-C. et al. A tecpr2 knockout mouse exhibits age-dependent neuroaxonal dystrophy associated with autophagosome accumulation. Autophagy 0, 1–14 (2020).

153. Buechler, C. et al. Expression pattern and raft association of NIPSNAP3 and NIPSNAP4, highly homologous proteins encoded by genes in close proximity to the ATP-binding cassette transporter A1. Genomics 83, 1116–1124 (2004).

154. Chander, P., Kennedy, M. J., Winckler, B. & Weick, J. P. Neuron-Specific Gene 2 (NSG2) Encodes an AMPA Receptor Interacting Protein That Modulates Excitatory Neurotransmission. eNeuro 6, (2019).

155. Ross, S. E. et al. Bhlhb5 and Prdm8 form a repressor complex involved in neuronal circuit assembly. Neuron 73, 292–303 (2012).

156. Inoue, M. et al. Prdm8 regulates the morphological transition at multipolar phase during neocortical development. PloS One 9, e86356 (2014).

157. VanGuilder Starkey, H. D. et al. Neuroglial expression of the MHCI pathway and PirB receptor is upregulated in the hippocampus with advanced aging. J. Mol. Neurosci. MN 48, 111–126 (2012).

158. Li, X.-H. et al. Gene Expression Profile of the Hippocampus of Rats Subjected to Chronic Immobilization Stress. PLoS ONE 8, e57621 (2013).

159. Stadhouders, R. et al. HBS1L-MYB intergenic variants modulate fetal hemoglobin via long-range MYB enhancers. J. Clin. Invest. 124, 1699–1710 (2014).

160. Baird, M. A., Hsu, T. Y., Wang, R., Juarez, B. & Zweifel, L. S. κ Opioid Receptor-Dynorphin Signaling in the Central Amygdala Regulates Conditioned Threat Discrimination and Anxiety. eNeuro 8, ENEURO.0370-20.2020 (2021).

161. Bodnar, R. J. Endogenous opiates and behavior: 2012. Peptides 50, 55–95 (2013).

162. Szklarczyk, K., Korostynski, M., Golda, S., Solecki, W. & Przewlocki, R. Genotype-dependent consequences of traumatic stress in four inbred mouse strains. Genes Brain Behav. 11, 977–985 (2012).

163. Johnson, M. A. et al. Chronic stress differentially alters mRNA expression of opioid peptides and receptors in the dorsal hippocampus of female and male rats. J. Comp. Neurol. 529, 2636–2657 (2021).

