## Supplemental Figures for "Transcriptomic profiles of stress susceptibility and resilience in the amygdala and hippocampus"

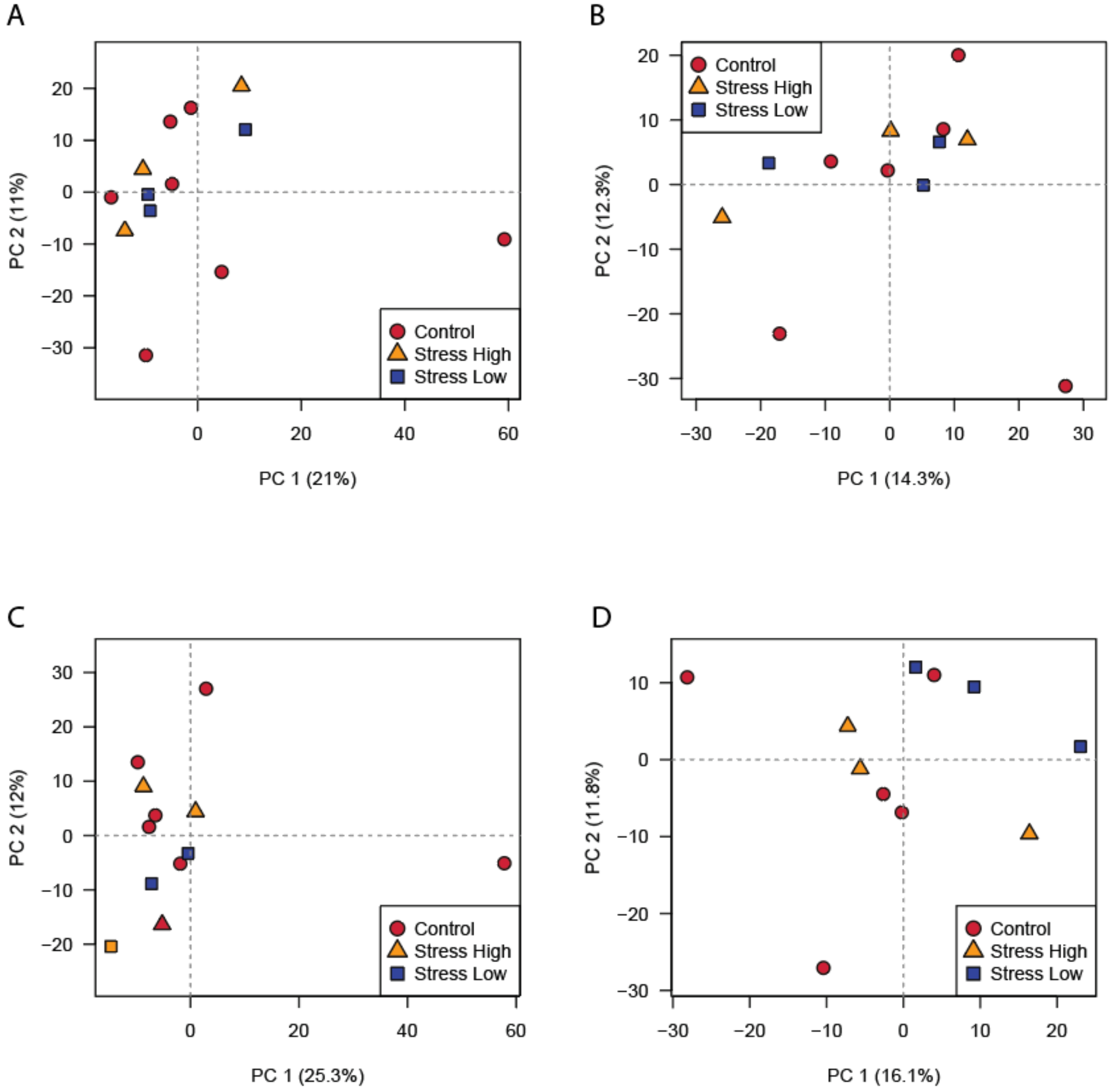


**Supplemental Figure 1:** PCA of normalized gene expression values for the amygdala and hippocampus dentate gyrus (DG) samples. A. Amygdala - all samples included. B. Amygdala - outlier control sample removed. C. DG - all samples. D. DG - outlier control sample removed.

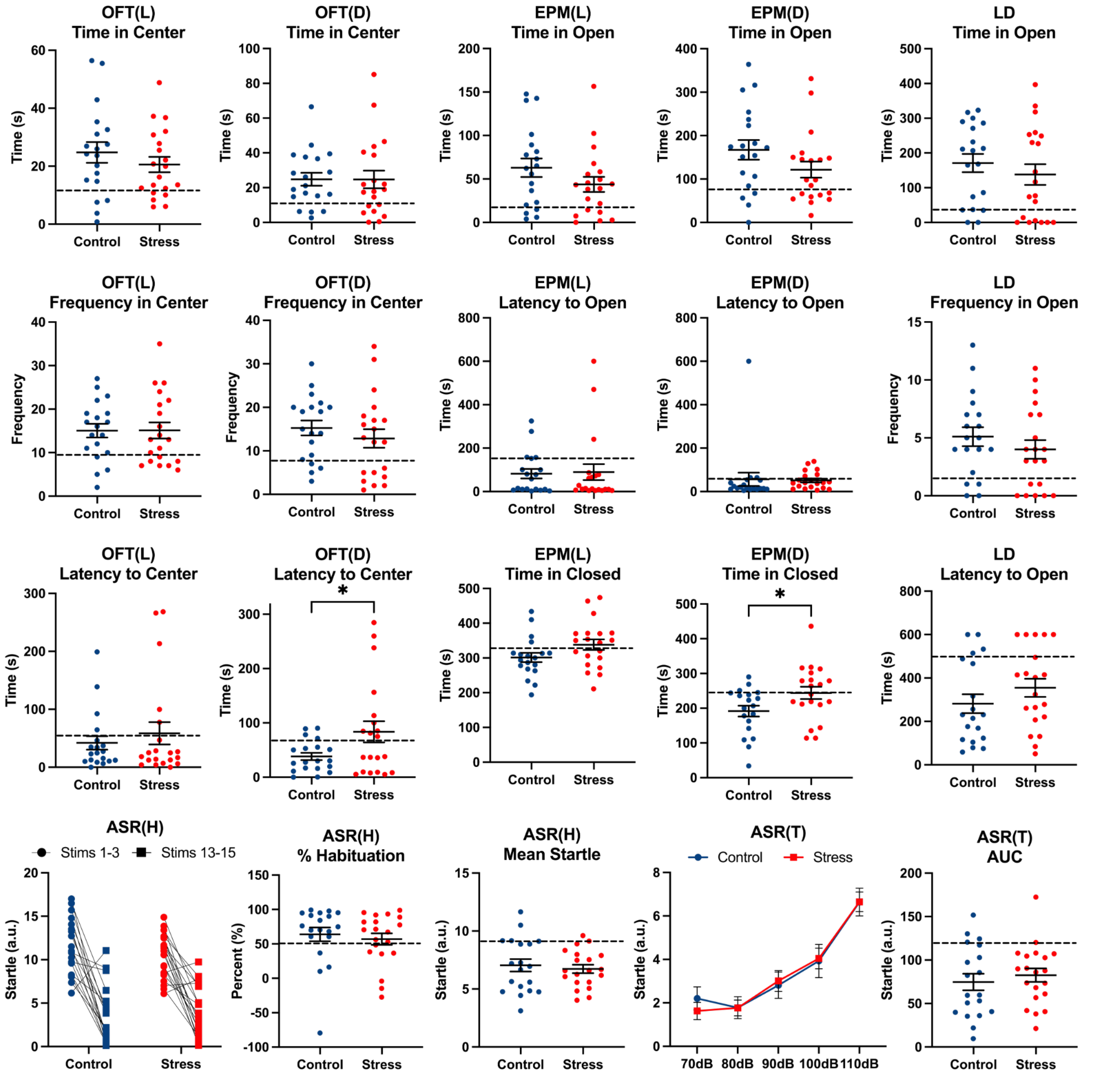


**Supplemental Figure 2:** All behavior data. Dotted line marks the 20th percentile of the control distribution. OFT(L), open field test under bright lights; OFT(D), open field test under dim lights; EPM(L), elevated plus maze under bright lights; EPM(D), elevated plus maze under dim red lights; ASR(H) 1-3, acoustic startle response test habituation phase, mean startle to stimuli 1-3; ASR(H) 13-15, ASR(H) mean startle to stimuli 13-15; ASR % Habituation, ASR(H) percent habituation; ASR(H), Mean all, mean startle response to all 15 stimuli; ASR(T) AUC, area under the curve from the ASR threshold phase. *$p < 0.05$

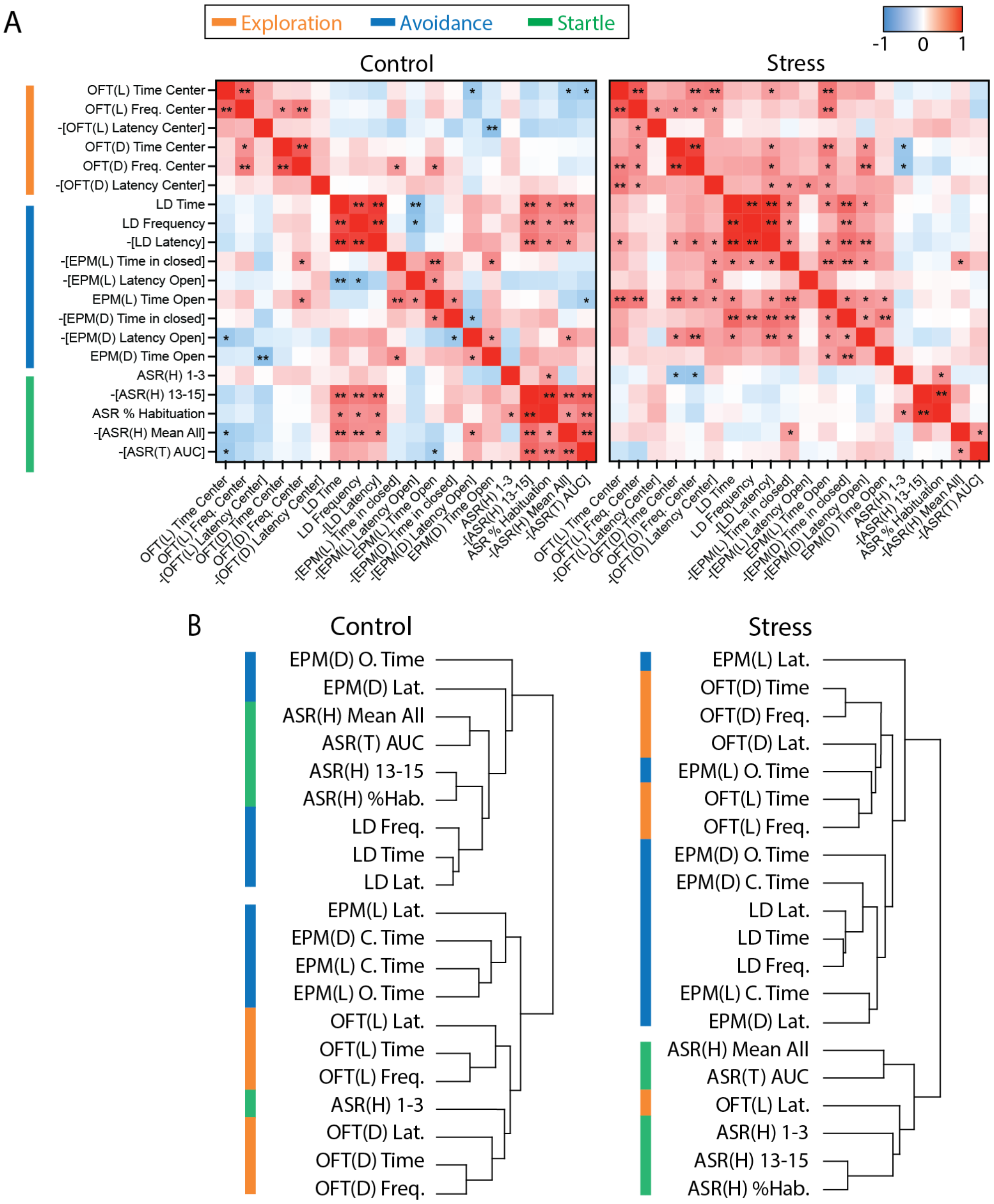


**Supplemental Figure 3:** Behavioral correlations and clustering. A. Heatmaps of correlations between behavioral metrics in control (left) and stress-exposed (right) rats. Pearson r values range from blue (-1) to red (+1), and asterisks within squares mark significant correlations. Colored bars classify individual behaviors according to their hypothesized representation in the literature (for example, elevated plus maze is typically used to measure anxiety-like avoidance behavior in rodents). For clarity of presentation, behavioral metrics with opposite valence (for example, latency and time spent in the closed arm of the elevated plus maze) from those of standard metrics (for example, time spent in the open arms of the elevated plus maze) were multiplied by -1. B. Hierarchical clustering of behavioral metrics in control (left) and stress-exposed (right) rats. OFT(L), open field test under bright lights; OFT(D), open field test under dim lights; EPM(L), elevated plus maze under bright lights; EPM(D), elevated plus maze under dim red lights; ASR(H) 1-3, acoustic startle response test habituation phase, mean startle to stimuli 1-3; ASR(H) 13-15, ASR(H) mean startle to stimuli 13-15; ASR % Habituation, ASR(H) percent habituation; ASR(H), Mean all, mean startle response to all 15 stimuli; ASR(T) AUC, area under the curve from the ASR threshold phase. *p < 0.05, **p < 0.01

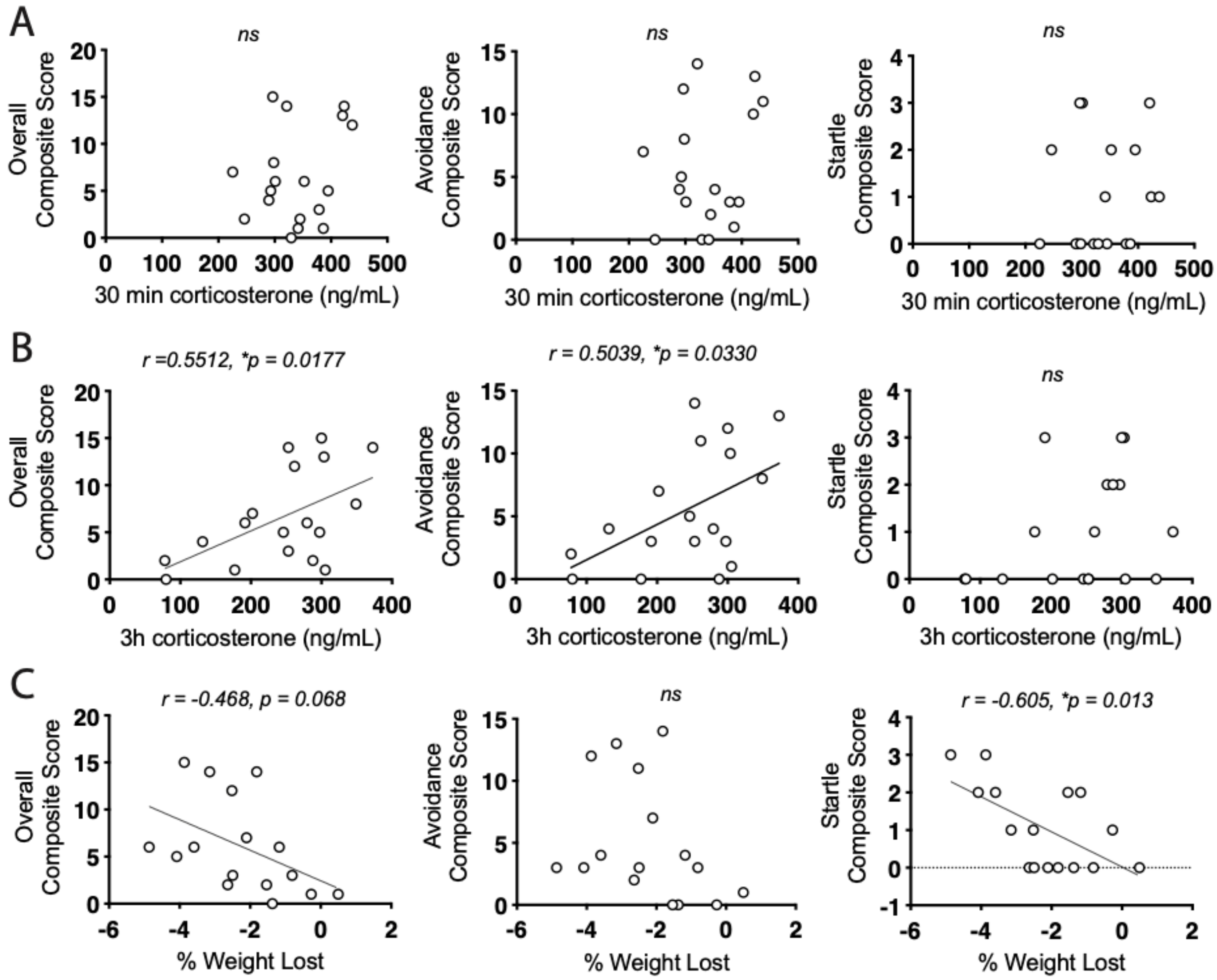


**Supplemental Figure 4:** Correlations between corticosterone, weight loss, and behavior after stress. A. Correlations between serum corticosterone at 30 minutes and overall (left), avoidance (middle), and startle (right) composite scores (overall: r = 0.2443, p = 0.3287; avoidance: r = 0.2263, p = 0.3665; startle: r = 0.1332, p = 0.5982). B. Correlations between serum corticosterone at 3 hours and overall (left), avoidance (middle), and startle (right) composite scores (overall: r = 0.5512, p = 0.0177; avoidance: r = 0.5039, p = 0.033; startle: r = 0.3275, p = 0.1846). C: Correlations between percent weight change from day of stress to day after stress and overall (left), avoidance (middle), and startle (right) composite scores (overall: r = -0.468, p = 0.068; avoidance: r = -0.3477, p = 0.187; startle: r = -0.605, p = 0.013). *p < 0.05

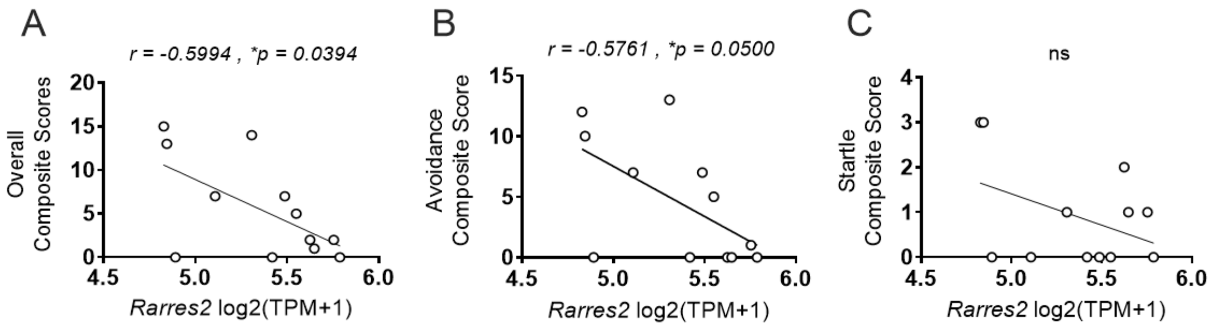


**Supplemental Figure 5**: *Rarres2* gene expression correlations to Overall Composite, Avoidance Composite and Startle Composite Score. A. Pearson correlation of *Rarres2* gene expression to Overall Composite Scores (r = -0.5994, p = 0.0394). B. Pearson correlation of *Rarres2* gene expression to Avoidance Composite Score (r = -0.5761, p = 0.0500). C. Pearson correlation of *Rarres2* gene expression to Startle Composite Score (ns).

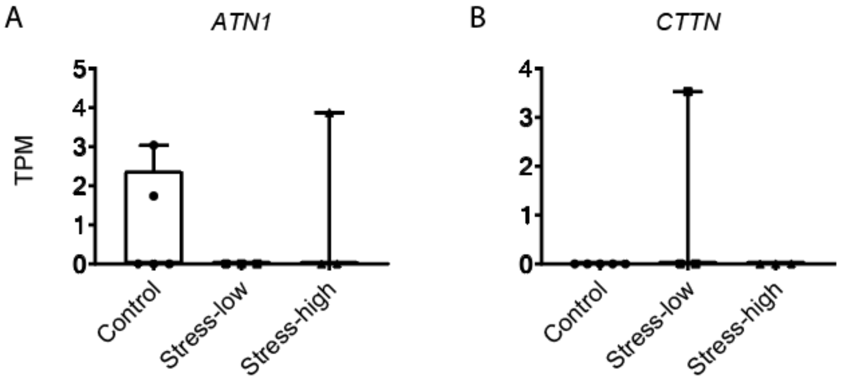


**Supplemental Figure 6:** *ATN1* and *CTTN* gene expression differences between control, stress-low and stress-high groups. A. *ATN1* gene expression (in TPM) for the 3 groups. B. *CTTN* gene expression (in TPM) for the 3 groups. Note that in both cases, outlier values drive group differences.

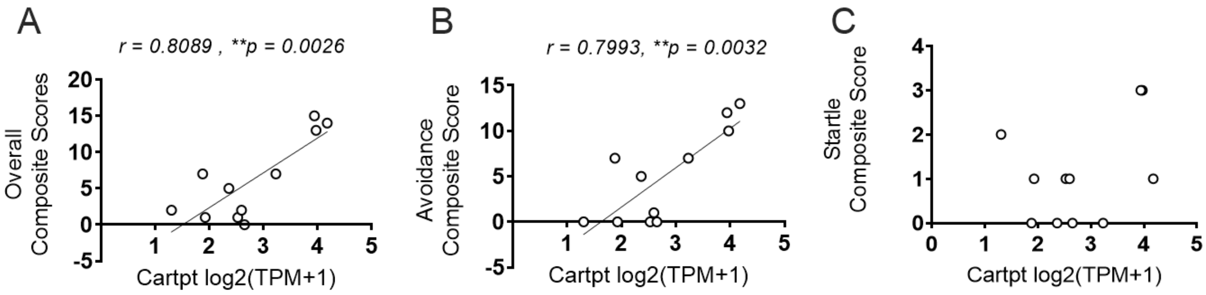


**Supplemental Figure 7:** *Cartpt* gene expression correlations to Overall Composite, Avoidance Composite and Startle Composite Score. A. Pearson correlation of *Cartpt* gene expression to Overall Composite Scores (r = 0.8089, p = 0.0026). B. Pearson correlation of *Cartpt* gene expression to Avoidance Composite Score (r = 0.7993, p = 0.0032). C. Pearson correlation of *Cartpt* gene expression to Startle Composite Score (ns).
